# Brain cycle flexibility and behavioral association

**DOI:** 10.64898/2026.11.22.768360

**Authors:** Quan Wang, Yubin Guo, Wanqing Nie, Qiushi Yang, Shuyi Zhang, Chongjing Luo, Yong Zhang, Xi-Nian Zuo

## Abstract

Canonical models of the macroscopic human connectome are dominated through static, cortico-centric networks by positive functional correlates. Yet naturalistic behavior demands fluid topological reconfigurations, wherein biologically critical functional antagonism (negative correlates) is routinely discarded as noise. Meanwhile, deep subcortical control remains overlooked. Here, we develop the Spatial Connectivity Cycle Flexibility (SCCF) framework to track whole-brain community dynamics across signed interactions. We show that canonical static networks reflect time-averaged projections of an underlying dynamic probability landscape, actively orchestrated by three (basal, regulatory, and fast) conserved subcortical routing modes. Individual dynamic modular co-occurrence function-prints predict 16 independent behavioral dimensions across cognitive, personality, and physiological domains, indicating that behavioral heterogeneity depends on the dynamic balance between network integration and antagonistic segregation. Crucially, independent cross-cohort validation reveals an organizational duality: subcortical routing for core cognitive computation is broadly conserved across cultures, whereas socio-emotional circuits exhibit marked divergence aligned with distinct behavioral ecologies. These findings challenge prevailing cortico-centric and positive-only assumptions of the functional connectome, providing an exploratory computational framework that links subcortical routing and active network antagonism to human phenotypic diversity and behavioral adaptation.

## Introduction

For decades, the macroscopic organization of the human brain has been characterized through the lens of functional segregation, establishing a canonical model of spatially stable intrinsic connectivity networks such as the default mode network (DMN) and frontoparietal network (FPN)[1, 2]. Designed primarily from time-averaged resting-state functional magnetic resonance imaging (fMRI), this framework captures the static modular organization of the brain. However, during naturalistic behavior, human cognition requires the brain to flexibly switch between internal reflection and external goal-oriented actions[3, 4]. To support these computational demands, high-resolution functional mapping and electrophysiological recordings indicate that macroscopic network boundaries are dynamically adjustable, with whole-brain functional coupling undergoing transient topological reconfigurations across rapid timescales[5, 6].

The paradigm shift toward dynamic functional connectivity (dFC) has provided important insights into these temporal fluctuations; however, prevailing models have largely focused on cortical reconfiguration and tracked macroscale dynamics primarily through positive temporal synchrony[7, 8]. Although computationally tractable, this focus places less emphasis on two complementary physiological mechanisms. First, physiological state transitions are accompanied by the reconfiguration of antagonistic functional networks[6, 9]. Specifically, the negative coupling (anticorrelation) between default mode and executive control networks systematically shifts between attentional states[10, 11]. In standard dFC pipelines that retain only positive weights, this competitive decoupling often remains unmodeled[12]. Second, large-scale changes in the cross-cortical state do not occur in isolation within the cortex but are closely modulated by deep subcortical projections[13, 14]. Beyond serving as afferent relays, subcortical structures (e.g., the thalamus and striatum) function as integrative routing hubs[15], operating through dense cortico-striato-thalamo-cortical loops to integrate behavioral goals[16, 17]. However, how these subcortical routing hubs dynamically coordinate the reconfiguration of antagonistic cortical modules remains largely unexplored in conventional dFC frameworks.

To examine how dynamic functional reconfiguration relates to static modular parcellations, here we develop the Spatial Connectivity Cycle Flexibility (SCCF) computational framework (Fig. 1). By extending traditional modularity detection algorithms[18, 19], we incorporate antagonistic negative connectivity as an explicit driver of dynamic network reconfiguration. Applying this framework to functional neuroimaging data indicates that cortical reorganization follows a three-tier hierarchy, while subcortical routing hubs exhibit marked functional flexibility[20] and contribute to macroscopic cortical reorganization through different routing modes. Building on Connectome-based Predictive Modeling (CPM)[21, 22], we further constructed a behavioral association framework using dynamic modular co-occurrence patterns (i.e. the frequency of nodal co-participation across functional modules) as predictors. This approach yielded 16 prediction targets including eight latent factors and eight independent components derived from 29 behavioral and physiological indices, which span cognitive abilities, personality and social traits, and physiological status. In particular, these associations depended on the retention of antagonistic (negative) connectivity along with positive coupling. We developed the framework in the Human Connectome Project (HCP; *N* = 1,031)[23], and then evaluated its cross-cohort consistency in an independent cohort of Han Chinese young adults with extensive behavioral measures (the I See Your Brain dataset, ISYB; *N* = 215). Together, these findings integrate signed network dynamics with subcortical routing, highlighting the role of subcortical structures in coordinating dynamic reconfiguration across the macroscopic brain.

**Fig. 1.**
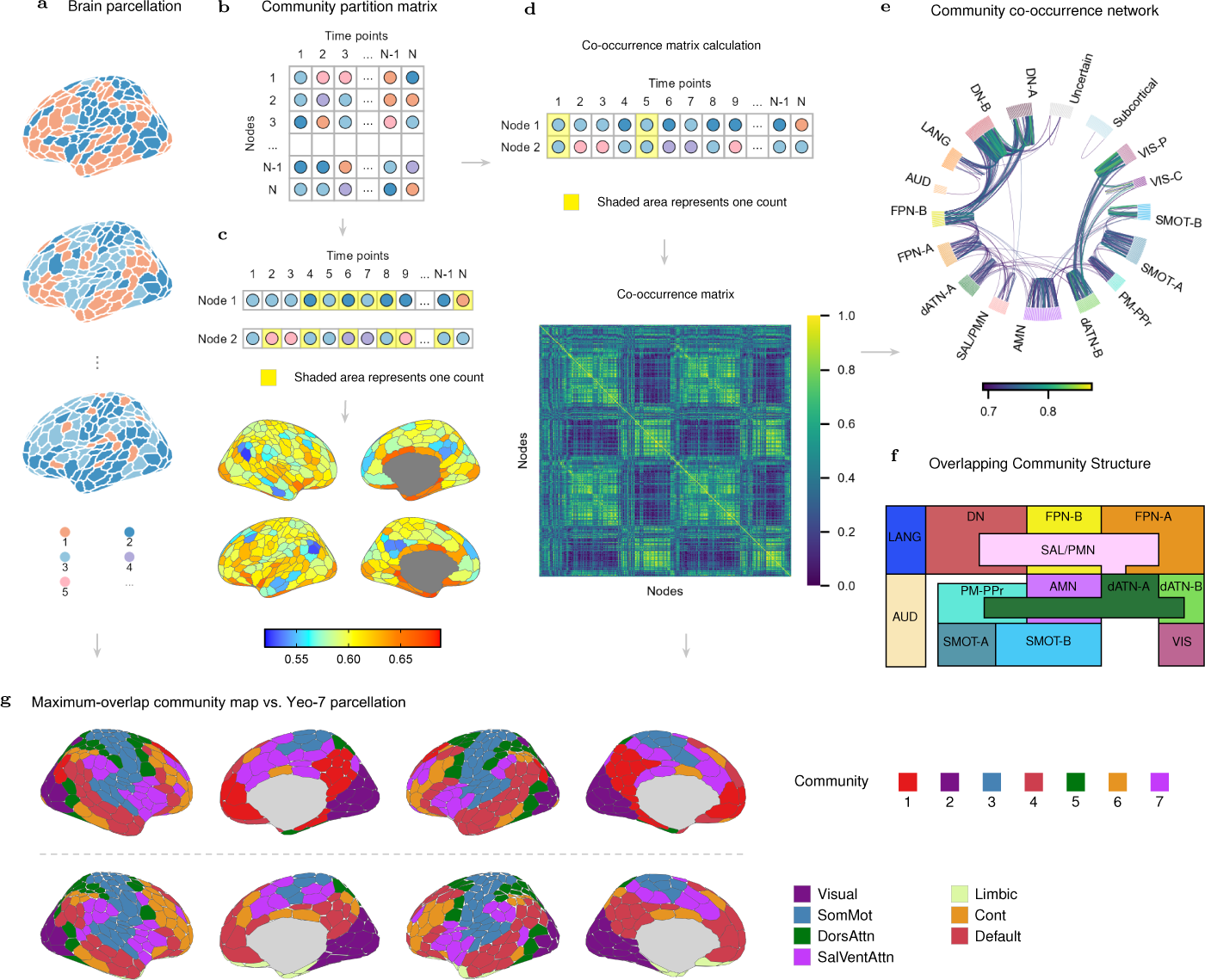
Spatial Connectivity Cycle Flexibility (SCCF) framework for mapping dynamic brain organization. **a**, Dynamic functional communities identified across time windows with a signed multilayer modularity-maximization algorithm; colours denote distinct communities. **b**, Community-affiliation matrix across different time windows, where rows represent distinct brain regions and columns represent different time windows. **c**, Spatial distribution of node flexibility, defined as the frequency at which a given brain region changes its community affiliation; colour bar, flexibility (0.52–0.69). **d**, Community co-occurrence matrix, representing the frequency at which each pair of brain regions is assigned to the same community. **e**, Network connectivity graph constructed from the predominant integration patterns of the community co-occurrence matrix, visualized with the Du-15 parcellation. **f**, Three-tier hierarchical organization derived from the predominant integration patterns in **e**: sensory systems (tier 3), attention/action systems (tier 2), and transmodal regulatory systems (tier 1). **g**, Maximum-overlap community map versus the Yeo-7 parcellation. Upper row, group-consensus communities obtained by the maximum overlap of the community co-occurrence matrix across time windows (arrow from **a**) and individuals (arrow from **d**) in the Human Connectome Project (HCP; *n* = 1,031); colours and numbers 1–7 denote distinct communities. Lower row, the Schaefer 400-parcel Yeo-7 parcellation, coloured by the seven Yeo networks: Visual, Somatomotor, Dorsal Attention, Salience/Ventral Attention, Limbic, Control, and Default.

### A hierarchical architecture of network flexibility

Conventional analysis of dynamic brain networks has focused primarily on temporal fluctuations in pairwise connectivity, with comparatively limited attention to how network modules reconfigure over time. Moreover, prevailing pipelines typically restrict network representations to thresholded, sparse matrices of exclusively positive connections. Such simplifications obscure signed physiological interactions, most notably the antagonism between the default mode (DMN) and action mode networks (AMN)[9, 24, 25], as well as higher-order topological organization driven by connectivity homophily[26, 27].

To overcome these methodological constraints, we developed the Spatial Connectivity Cycle Flexibility (SCCF) framework to track community dynamics across both positive and negative functional interactions. Specifically, SCCF constructs a multilayer network from full-band functional-connectivity affinities[21, 28, 29] and applies a generalized multilayer modularity-maximization algorithm[19, 20] that incorporates signed edge weights[22, 30] (see Appendix A). By preserving both the sign and strength of interregional relationships, this formulation enables the tracking of time-varying modular reconfigurations, providing a principled foundation to map their spatial distribution across the cortex.

At the cortical topographic level, the SCCF framework reveals a dynamic reorganization landscape that departs from traditional static boundaries. Highly flexible regions are predominantly located along topological borders where multiple functional modules intersect (Fig. 1c; Appendix B.1). For example, perisylvian cortical areas, positioned at the anatomical crossroads of auditory, somatomotor, language, action mode, and default mode networks[31], exhibit pronounced dynamic flexibility. In contrast, regions with minimal flexibility are restricted to topological cores that maintain stable modular affiliations over time, exemplified by the angular gyrus and the posterior cingulate cortex (PCC) remaining anchored to the DMN core.

Beyond identifying where reconfiguration occurs in space, tracking the temporal modular co-occurrence among these regions uncovers a fundamental organizing principle: a “three-tier hierarchical reorganization motif” that bridges local specificity with macroscopic network cycles (Fig. 1f; Appendix B.2). On a broader scale, this cyclical reorganization aligns with the system-level evolutionary and sensory-fugal cascade of the brain [31–33]. Along this processing trajectory (Fig. 1f), unimodal primary sensory networks (tier 3) maintain intrinsic cohesion; intermediate attention and action networks (tier 2), including dorsal attention networks (dATN-A and dATN-B), premotor and action mode networks, undergo frequent modular switching to serve as dynamic routing relays; and transmodal regulatory networks (tier 1; notably FPN and DMN) operate as convergence hubs for higher-order synthesis.

Rather than operating as a rigidly sequential, strictly feedforward hierarchy, this architecture supports flexible cross-tier interactions essential for complex cognition. First, at the sensory-to-transmodal interface, the primary auditory network bypasses intermediate relays, leveraging high-frequency modular co-occurrences to establish a direct “cross-tier bypass” with DMN-associated language areas. This direct functional bridge reflects the computational specialization of auditory-linguistic processing and provides macroscopic dynamic evidence for an accelerated auditory-language axis[34, 35]. Second, at the transmodal apex (tier 1), the DMN does not act as a passive informational sink[32]. Although predominantly integrating with FPN-B, the DMN also selectively recruits the salience network (SAL/PMN) as a transient intermediary for “internal feedback routing.” This selective recruitment dynamically calibrates the balance between cooperation and antagonism across frontoparietal control systems[36, 37], thereby explaining the increased topological flexibility observed in SAL / PMN regions. Finally, in the intermediate attention level (tier 2), dATN-A acts as a cross-network nexus that bridges PM-PPr, AMN and dATN-B, operating in concert with SAL/PMN to facilitate integrative information routing (Appendix B.2).

These fluid moment-to-moment reconfigurations raise the fundamental question: how does such pervasive dynamic flexibility reconcile with canonical, static functional networks? In our framework, aggregating the most frequent dynamic modular co-occurrences (maximal overlap) across time and individuals naturally recovers classical, time-averaged network parcellations[2, 31] (Appendix B.3). Rather than acting as fixed anatomical compartments, canonical static networks simply represent the most probable time-averaged configurations accumulated from ongoing dynamic interactions[38, 39]. Crucially, although this dynamic hierarchical landscape unfolds throughout the cerebral cortex, our spatial mapping reveals that large-scale cortical switching does not occur in cortical isolation; instead, it is closely coupled and supported by underlying subcortical dynamics[17, 40].

### Three subcortical modes for cortical reconfiguration

Although the SCCF framework reveals hierarchical reorganization within the cerebral cortex, spatial mapping indicates that cortical topological switching is restricted mainly to localized boundaries between functional networks. By contrast, subcortical regions exhibit transitions that span multiple macroscopic networks. This difference in spatial scale suggests that the reconfiguration of the functional connectome is driven primarily by the subcortex[13, 14, 17]. Mapping the dynamic modular co-occurrence profiles of subcortical nodes showed that these structures form a functional switching network that orchestrates transitions across diverse brain systems. Profiling subcortical-cortical integration trajectories across all subcortical nodes yielded three conserved routing modes that drive macroscopic cortical reconfiguration (Fig. 2; Appendix B.4): the Basal Mode, the Regulatory Mode, and the Fast Mode.

**Fig. 2.**
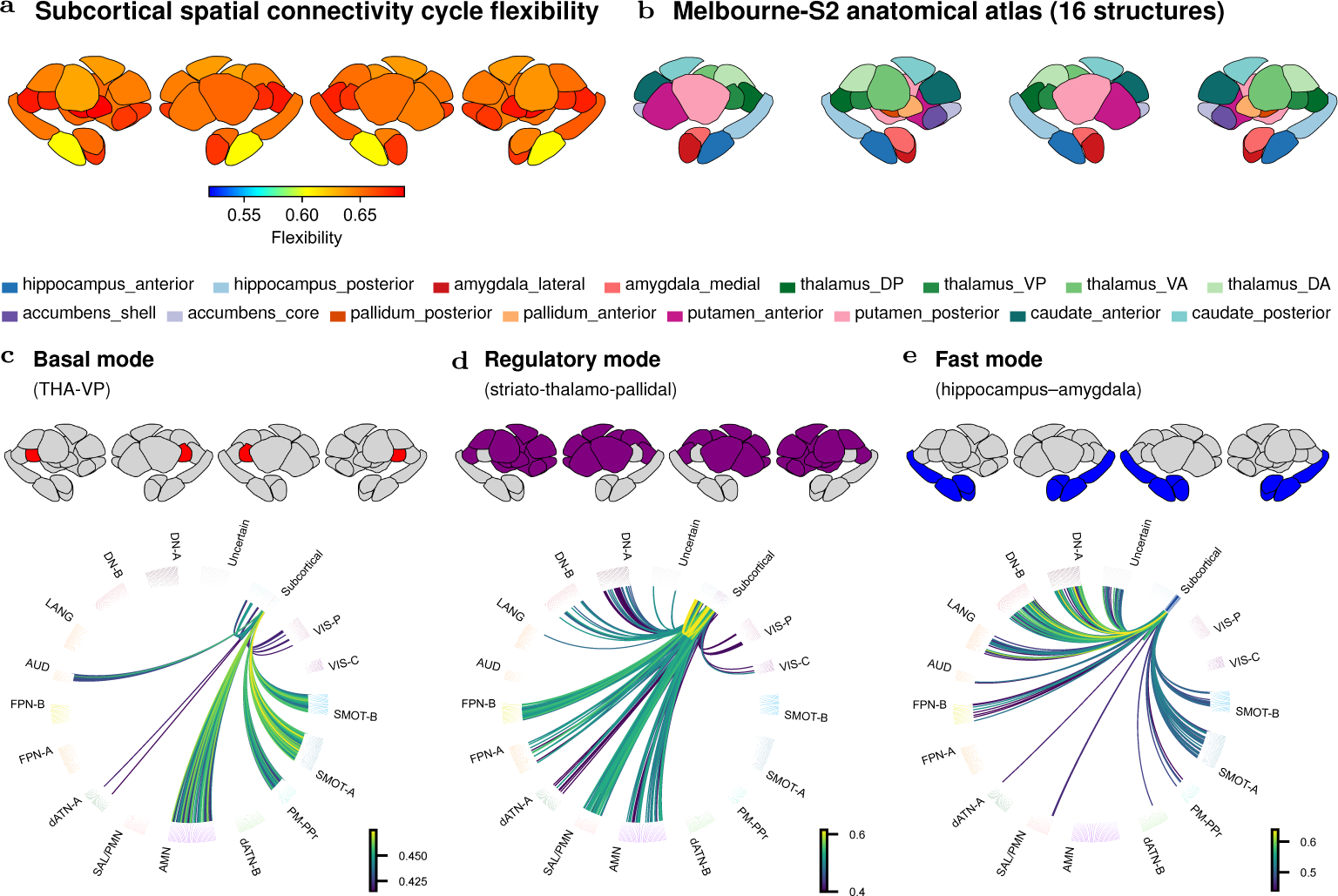
Spatial organization of subcortical connectivity cycle flexibility and three functional routing modes. **a,** Spatial distribution of Spatial Connectivity Cycle Flexibility (SCCF) across subcortical parcels in the Human Connectome Project (HCP; *n* = 1,031), displayed from four views of the subcortical surface. Colours indicate SCCF values (0.52–0.69). **b,** Anatomical parcellation of subcortical structures according to the Tian (Melbourne-S2) atlas, shown in the same four views. The 16 subcortical structures are colour-coded by nucleus group, as indicated in the legend below. **c–e,** Three subcortical routing modes characterized by distinct patterns of dynamic subcortical–cortical integration. For each mode, the upper panels highlight the associated subcortical parcels across four anatomical views, while the lower connectogram illustrates the frequency with which subcortical parcels and cortical regions belonging to different functional networks are integrated into the same dynamic module. The colour scale indicates the corresponding co-occurrence frequency. **c, Basal Mode.**Centred on the ventral posterior thalamus (THA-VP), this mode predominantly connects primary sensory–motor networks, including auditory, motor and visual networks, with intermediate attention-related networks, particularly the action-mode and premotor networks. **d, Regulatory Mode.** Distributed across the associative thalamus, striatum and globus pallidus, this mode coordinates connections among the default mode (DMN), frontoparietal (FPN) and action-mode (AMN) networks. The associative thalamus, nucleus accumbens and caudate connect all three networks, whereas the putamen and globus pallidus predominantly link the FPN and AMN. **e, Fast Mode.**Centred on the hippocampus and amygdala, this mode is characterized by direct cross-tier connections between higher-order transmodal networks (tier 1) and primary sensorimotor networks (tier 3), bypassing intermediate attention-related networks (tier 2).

The Basal Mode (Fig. 2c) centers on the ventral posterior thalamus (THA-VP). At the macroscopic network level, this mode is characterized by dense functional integration between primary sensory-motor networks (the auditory, motor, and visual networks) and intermediate attentional-tier networks (primarily the action mode and premotor networks). Although classical anatomical frameworks assign the THA-VP primarily to a somatosensory relay[13, 41], its dynamic community co-occurrence reveals a broader system-level role: it provides the substrate for continuous bottom-up information transfer. By coupling perceptual inputs with attentional prioritization, the THA-VP acts as a multisensory integrator[15], maintaining basic perception and somatic vigilance while higher-order transmodal networks reconfigure[6].

In contrast, the Regulatory Mode (Fig. 2d) functions as the brain’s overarching cognitive switcher. Supported by a widespread cortico-striato-thalamic circuit (spanning the associative thalamus, striatum, and globus pallidus), it coordinates dynamic coupling among the default mode (DMN), frontoparietal (FPN) and action-mode (AMN) networks. At the regulatory end of this cascade, the associative thalamus[14], nucleus accumbens[42], and caudate[43] bridge all three networks, integrating internal motivation with goal-directed cognitive strategies. At the end of execution, the putamen and globus pallidus couple the FPN with the AMN to mediate automated sensorimotor habits[44] and gate behavior output[45], working in concert with the regulatory end to translate cognitive regulation into external execution. Ultimately, by recruiting coordinated interactions across multiple networks that span regulatory and attention tiers, this mode enables the brain to flexibly adapt to complex and rapidly changing external environments.

Finally, the Fast Mode (Fig. 2e), anchored by the hippocampus and amygdala, establishes a non-hierarchical information-processing pathway by directly coupling tier-1 (e.g. DMN) and tier-3 (primary sensorimotor) modules, thereby bypassing the intermediate tier-2 systems responsible for attention and information transmission. Consistent with the evolutionarily conserved role of the limbic system in affective and mnemonic processing[46], this cross-tier architecture serves as an evolutionary fast track: it channels external sensory input directly into internal affective and memory systems and promptly translate them into immediate motor responses during environmental crises[47]. Beyond acute survival responses, this limbic-cortical coupling also shapes individual differences in emotional reactivity, social functioning, and personality traits[48]. Consequently, the Fast Mode provides an affect-driven shortcut for rapid global network reorganization.

Collectively, the Basal, Regulatory, and Fast routing modes constitute an evolutionary division of labor within the subcortex: the Basal Mode sustains sensory-vigilance grounding, the Regulatory Mode supports goal-directed cognitive switching, and the Fast Mode provides an affect-driven bypass for rapid environmental adaptation. Together, these complementary mechanisms indicate that subcortical structures act as physiological engines driving macroscopic cortical reconfiguration. Crucially, because these hubs set the dynamic balance between global integration and segregation, we hypothesized that individual variations in these routing trajectories could decode differences in complex cognitive and behavioral phenotypes, helping to establish brain-behavior associations that have remained elusive[49].

### SCCF for Brain-Wide Association Studies

The continuous reorganization of brain networks is more than a mere physiological oscillation; it is an evolved computational strategy to adapt to complex environments[4, 6]. To test whether our SCCF architecture can decode human phenotypic diversity, we rely on connectome-based predictive modeling[21] to convert dynamic modular co-occurrence patterns into individual dynamic modular co-occurrence fingerprints (DMCFs). These DMCFs achieved a cross-run individual identification accuracy of 62.49% via nearest-neighbor matching (chance level 0.08%) and were further compressed into behavior-specific frequency signatures (Appendix A.18). In the HCP cohort, we evaluated these signatures in 29 behavioral and physiological indices, which were summarized into 16 target phenotypes (eight latent factors and eight independent components) spanning three core domains[50] (Fig. 3a; Appendix B.5). Across these three domains, DMCF signatures significantly predicted higher-order cognitive abilities (e.g. verbal intelligence, *r* = 0.457*, P <* 0.001), personality and socio-emotional traits (e.g. extraversion and friendship, *r* = 0.11; social distress, *r* = 0.11) and basic physiological indicators (e.g. sleep quality, *r* = 0.19; blood pressure, *r* = 0.18; Fig. 3b). Consequently, spatial cyclic flexibility serves a dual role: providing the computational substrate for higher-order cognition while remaining tightly coupled to physiological homeostasis[51, 52].

**Fig. 3.**
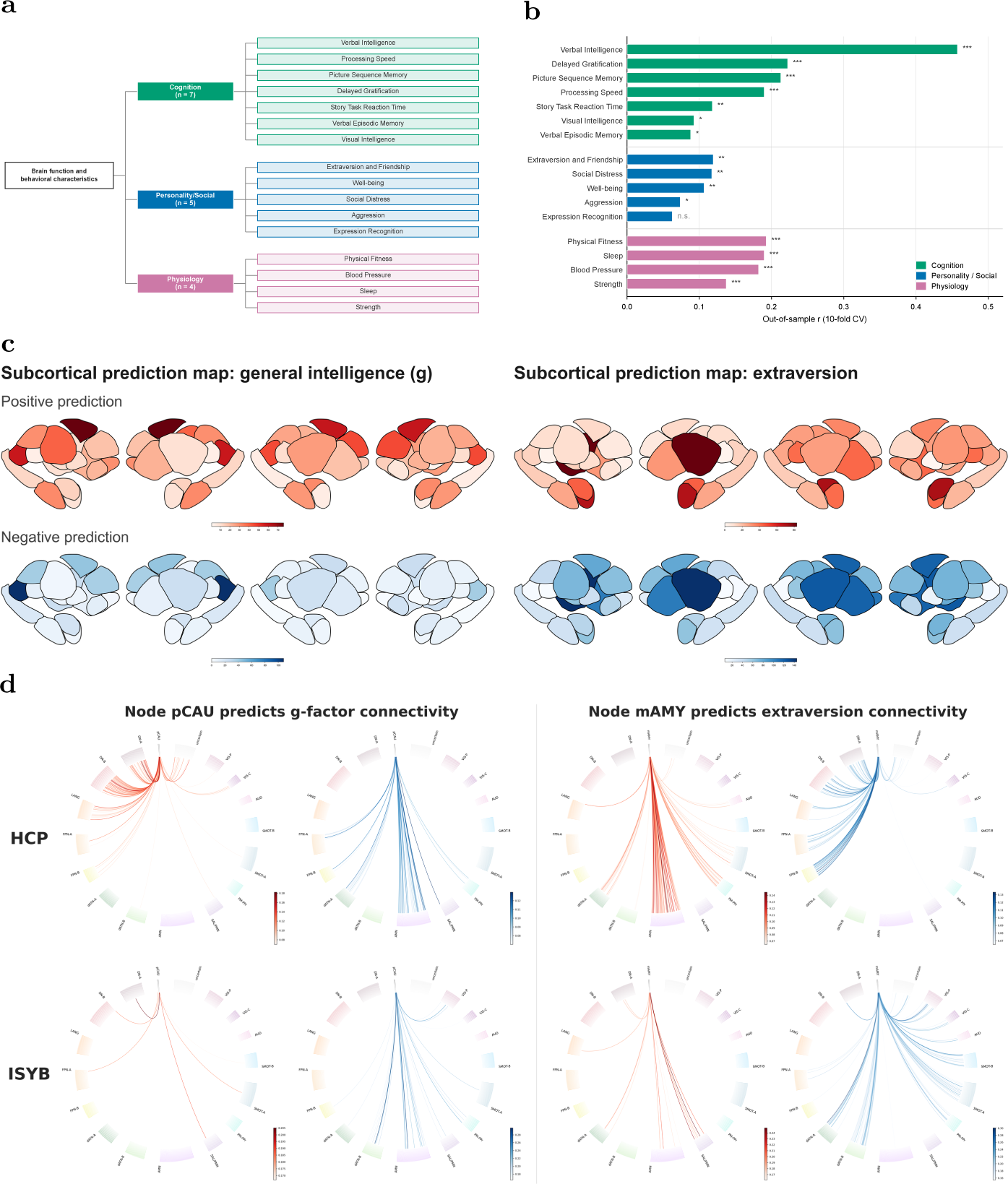
Phenotypic prediction and cross-cohort subcortical routing of dynamic modular co-occurrence fingerprints (DMCFs). **a**, Hierarchical taxonomy of the 16 HCP target phenotypes across cognitive (green), personality/social (blue), and physiological (magenta) domains, summarized from 29 indices into eight latent factors and eight independent components. **b**, Out-of-sample prediction performance from full DMCFs (10-fold cross-validated kernel ridge regression, pooled Pearson *r*; *n* = 828 for latent factors; *n* = 991–1,031 for independent components). Asterisks indicate significance surviving within-family Benjamini–Hochberg FDR correction (eight latent factors and eight independent components corrected separately; *q*_BH_ *<* 0.05; one-sided permutation *^∗^P <* 0.05, *^∗∗^P <* 0.01, *^∗∗∗^P <* 0.001; n.s., not significant; Appendix A.20). **c**, Subcortical predictive degree maps in the HCP (4-run subset) for general intelligence (*g*, left; *n* = 785) and extraversion (right; *n* = 917). Shading reflects the number of uncorrected *P <* 0.05 edges per subcortical parcel, shown separately for positive (integration, red) and negative (segregation, blue) predictions. **d**, Cross-cohort connectograms comparing HCP (top) and I See Your Brain (ISYB, bottom) cohorts for cognitive routing (posterior caudate [pCAU] predicting *g* in HCP and intelligence factor 2 in ISYB; left) and socio-emotional routing (medial amygdala [mAMY] predicting extraversion; right). Positive (red, left ring) and negative (blue, right ring) subcortical–cortical edges are shown separately; colour bars indicate edge strength.

As the hubs driving global network reconfiguration (Fig. 2), subcortical structures encoded intelligence, personality, and bodily-control phenotypes by combining the three routing motifs described above (Fig. 3b–d). First, in the cognitive and action-control hubs, comprising the posterior caudate (pCAU), dorsoposterior thalamus (THA-DP) and posterior putamen (pPUT), pCAU and THA-DP supported core cognitive computations[14]. The pCAU operated as a functional switcher between the default mode (DMN) and the action mode (AMN) networks[16, 53, 54]: its integration with the DMN was associated with verbal intelligence and agreeableness, reflecting a shared substrate for language-based cognition and pro-social conformity, while its segregation from the AMN supported visual intelligence. Within the same system, pPUT acted as an executive motor gate for social action[55, 56]. Tighter integration with first-tier (higher-order regulatory) networks, together with stronger segregation from second- and third-tier (attentional and sensorimotor) networks, was associated with greater extraversion, indicating that extraverted individuals translate internal intentions into overt social actions with reduced behavioral inhibition[57]. Second, in socio-emotional and experiential hubs, both the medial amygdala (mAMY) and anterior hippocampus (aHIP) shaped personality traits by decoupling from first-tier regulatory networks while integrating with external perceptual and action systems. Specifically, sparser coupling of the mAMY with first-tier networks, combined with tighter coupling to the AMN and sensorimotor networks, was associated with greater extraversion[58, 59], indicating that affective responses in extraverts depend more on direct external sensory interaction and action engagement than on internal cognitive processing. Similarly, a greater co-occurrence of aHIP with second and third level communities, together with a reduced co-occurrence with first level communities, was associated with higher trait openness[60, 61], meaning that open individuals incorporate broader external sensory material to fuel constructive imagination while experiencing less top-down control and inhibition from first-tier networks (Appendix B.7).

To test the cross-population generalizability of the DMCF architecture, we applied the framework to an independent East Asian cohort (“I See Your Brain”, ISYB; *N* = 215 for imaging and *N* = 162 for behavioral analyzes), revealing a clear cross-cultural principle: the routing topology of cognitive computation was conserved, whereas representations of socio-emotional personality diverged (Fig. 3d; Appendix B.9). In the cognitive domain, intelligence-related edges of the pCAU and THA-DP identified in ISYB also tracked general intelligence (*g*; the general factor of ten HCP cognitive measures[62]; Appendix A) in the Western cohort (HCP); meanwhile, the cross-dataset consistent pCAU agreeableness network and aHIP openness network predicted *g* and visual intelligence in HCP, respectively, indicating that the cross-culturally conserved aspects of personality traits are their embedded cognitive components[57]. By contrast, extraversion showed strong cultural specificity, with most of the significant shared edges showing opposite signs. In HCP, high extraversion was driven by mAMY–AMN integration, reflecting active external search, whereas in ISYB it relied on mAMY segregation from dorsal attention networks (dATN-A and dATN-B)—a release from internal cognitive monitoring consistent with collectivist norms of behavioral restraint[63, 64]. Notably, despite this pathway divergence, extraversion, and agreeability converged onto a shared mAMY routing substrate that predicted perceived social support in ISYB and the extraverted friendship factor in HCP. The core cognitive routing therefore forms a computational backbone that is conserved cross-culturally, while subcortical socio-emotional circuits tailor their routing strategies to cultural ecologies and converge to support human social affiliation[65, 66].

## Discussion

For decades, the macroscopic organization of the human brain has predominantly been conceptualized through static, cortico-centric frameworks [67, 68]. By developing the SCCF framework, this study moves beyond these conventional boundaries, indicating that canonical “static” functional atlases primarily reflect time-averaged projections (or metastable statistical baselines) of an underlying, highly dynamic probability landscape [38]. Although dynamic connectomics has increasingly captured temporal reconfigurations [38, 39], existing paradigms have largely overlooked the subcortex, continuing to focus on cortico-cortical interactions [40, 68]. Our findings challenge this persistent cortico-centric bias. By mapping global topological transitions, we show that subcortical structures (including the thalamus, striatum, hippocampus, and amygdala) act not only as relay stations, but as routing hubs that orchestrate large-scale cortical reconfigurations [40], fluidly coordinating transitions between internally guided cognition and externally focused action [69].

Beyond positive correlations among blood-oxygen-level-dependent (BOLD) signals, our framework establishes that functional antagonism captured through negative correlations constitutes an indispensable computational dimension for decoding complex human phenotypes. Although conventional network pipelines frequently discard negative couplings as methodological artifacts or threshold them out during graph construction [12, 30], our findings indicate that anti-correlated dynamics carry behaviorally relevant information [9, 10]. Empirically, within dynamic network reconfigurations, individuals exhibiting a reduced frequency of co-assigning the posterior caudate and action-mode networks to the same functional module displayed superior general intelligence, a modular segregation fundamentally rooted in negative signal correlations [70]. Intriguingly, in linking these subcortical reconfigurations to behavior, we observed pronounced functional lateralization across both intelligence and personality profiles, suggesting that subcortical routing hubs partition their computational capacity to support asymmetric cognitive and affective demands. Although the exact biological underpinnings of these anti-correlations warrant future validation, several non-mutually exclusive mechanisms may account for these observations. Negative correlations may reflect active inhibitory-mediated signal transmission [71, 72]; alternatively, persistent antagonism between discordant nodes may introduce computational interference if integrated into the same community, such that modular segregation minimizes functional cross-talk [73]; or such antagonism may foster functional specialization, insulating regions to dedicate local neural resources to domain-specific computations [73, 74].

The capacity of our framework to decode widespread brain–behavior associations suggests that it captures a fundamental organizational principle of the human connectome: the brain realizes complex function through the dynamic, flexible orchestration of multi-scale network integration [6, 74]. Rather than linking phenotypes directly to conventional static edge-level functional connectivity (a strategy prone to spatial-topography confounds and small effect sizes in phenotype prediction [49, 75]), our method quantifies the temporal frequency with which individual nodes are assigned to specific functional modules [20, 76]. By compressing high-dimensional network dynamics into discrete modular allegiance frequencies, this approach reveals that cognitive and behavioral capacity emerges from the system’s ability to maintain a functional repertoire of integration states [77]. In this view, individual differences in phenotypic diversity reflect stable biases in how routing hubs reconfigure their modular allegiance to navigate between segregation (specialized processing) and integration (global synthesis) over time [50, 51]. Consequently, the SCCF framework yields a tractable, low-dimensional representation that facilitates the reverse engineering of brain function. Coupled with kernel ridge regression, these compact topological features robustly predicted general intelligence as well as personality and social dimensions, demonstrating that low-dimensional modular dynamics effectively isolates behaviorally relevant individual differences to illuminate the neural basis of complex behavior [21, 78]. Despite these insights, the exact generative mechanisms underlying these dynamic integration states, and specifically how they evolve over time?, remain to be fully elucidated [39, 79]. Future work is critical to determine whether these modular allegiance patterns represent stochastic fluctuations or deterministic state-transition trajectories [80, 81], and how they interact with exogenous demands across varying temporal scales. Uncovering the time-varying nature of this organizational principle will be essential to understanding how the human brain maintains robust functional outcomes amidst the inherently noisy and non-stationary landscape of cognitive computation [77, 79].

Methodologically, our framework refines multilayer modularity maximization [19] through continuous-time Markov stability, modeling flow-based topological information propagation through intrinsic Markov diffusion time *t* and interlayer coupling matrix *C* [82, 83]. Although this formulation yields behaviorally predictive partitions, three methodological dimensions warrant systematic exploration to map the optimal subspace for brain–behavior alignment. First, within the parameter space, our preliminary findings indicate that whereas *C* primarily enforces temporal continuity across adjacent time slices with minimal impact on downstream phenotypic associations, Markov time *t* governs hierarchical topological resolution [84] and critically modulates predictive power. Second, the sliding–window duration - conventionally chosen via empirical heuristics [38, 39]—likely exhibits phenotype-specific optimality, as the window length dictates the cutoff frequency for infraslow oscillations [85] and distinct cognitive or behavioral domains operate across specialized BOLD frequency spectra [86, 87]. Finally, because our current algorithm represents a first-order Markovian approximation, extending it to second- or higher-order random walks [88, 89] is needed to capture non-Markovian neural trajectories; such higher-order diffusion processes preserve finer dynamical fidelity closely linked to shorter-period (higher-frequency) BOLD bands. Integrating systematic parameter profiling, frequency-resolved temporal windowing, and higher-order walk extensions will thus be pivotal for advancing reverse engineering of human brain function.

Despite capturing system-level topological principles, our framework highlights a critical frontier: the necessity of mapping modular reconfiguration within specific frequency bands. Increasing evidence suggests that brain networks do not operate as a monolithic entity, but rather as a multiplex system where different functional modules emerge and dissolve at different frequency scales [90, 91]. For example, infraslow oscillations may reflect the slow-evolving constraints of physiological homeostasis and global state transitions, while higher-frequency bands likely carry the rapid and task-specific computations required for flexible cognition [86, 87, 92]. We hypothesize that the modular reconfigurations observed here are not uniform across frequencies but are frequency-dependent signatures of specialized computational processes. Consequently, a comprehensive reverse engineering of brain function requires a shift from wide-band to frequency-resolved modular analysis, enabling us to isolate which specific BOLD spectral signatures track distinct behavioral or physiological phenotypes. Empirically validating this frequency-dependent organizational principle will be essential to understanding how the human brain segregates and integrates information across multiple temporal channels, thereby transforming our understanding of the connectome from a static or broad-band representation into a sophisticated, frequency-tuned computational architecture.

In summary, by establishing the subcortex as the active orchestrator of dynamic integration and antagonism across the human connectome, our findings bridge subcortical sub-architectures with macroscale network transitions. Across two culturally distinct cohorts, this dynamic coordination is captured by a subcortically driven tri-motif interaction model that decodes 16 independent cognitive, personality, and physiological components, offering a unified computational framework for human phenotypic diversity. Beyond identifying candidate topological targets for precision neuromodulation in psychiatric and neurodevelopmental disorders, this model provides a new perspective for investigating the dynamic organization of the human brain.

## Acknowledgements

We thank all the computational and logistical support provided by the National Basic Science Data Center “Chinese Data-sharing Warehouse for In-vivo Imaging Brain”. The authors also thank Peng Gao for assisting with the preprocessing of the ISYB data set.

## Declarations

### Funding

Scientific and technological innovation 2030 - the major project of the Brain Science and Brain-Inspired Intelligence Technology, 2021ZD0200500

### Conflict of interest/Competing interests

The authors have declared no competing interests.

### Ethical statement

All participants provided written informed consent. All the protocols were approved by the Institutional Review Boards at Institute of Psychology, Chinese Academy of Science (ISYB) and Washington University in Saint Louis (HCP).

### Data and code availability

Data were provided [in part] by the Human Connectome Project (HCP), WU-Minn Consortium (Principal Investigators: David Van Essen and Kamil Ugurbil; 1U54MH091657). ISYB data are deposited in the Chinese Color Nest Data Community (CCNDC) in the Science Data Bank (https://cstr.cn/31253.11.sciencedb.00740). All codes are publicly accessible at CCNDC (https://cstr.cn/31253.11.sciencedb.o00133.00031).

### Author contribution

QW: Conceptualization, Data curation, Formal analysis, Software, Visualization, Writing – original draft, review & editing. YG: Data curation, Investigation, Writing – original draft. WN: Investigation, Writing – original draft. QY: Investigation, Writing – original draft. SZ: Formal analysis, Investigation, Writing – original draft. CL: Validation, Investigation, Writing – review & editing. YZ: Supervision, Writing – review & editing. X-NZ: Conceptualization, Formal analysis, Software, Visualization, Funding acquisition, Supervision, Writing – review & editing.

## Notes

https://www.humanconnectome.org/study/hcp-young-adult/data-releases

https://doi.org/10.11922/sciencedb.00740

### Summary of Updates

Update of Affiliation for Yong Zhang. Revise the subsection 'Data and code avaliablity'.

