## Supplementary Materials for "Brain cycle flexibility and behavioral association"

### Appendix A Supplementary Methods

#### A.1 Participants and Datasets

Participants were recruited from the Human Connectome Project (HCP) S1200 data release [23] and the Chinese Multimodal Neuroimaging Database I See Your Brain" (ISYB) [93].

For the HCP dataset, 1,031 unique healthy adults each had at least one BOLD run in the resting-state that passed the standard quality control pipeline of HCP [94]. Usable data were available for 1,028 participants in the REST1_LR run, 972 in REST1_RL, 925 in REST2_LR, and 922 in REST2_RL; 917 participants contributed to all four runs. Among the 1,031 participants, 558 were female and 473 were male, aged 22–37 years (mean age 28.75 years, standard deviation 3.70). No participant had a history of neurological or psychiatric disorders. According to the requirements of the Institutional Review Board of Washington University in St. Louis, written informed consent was obtained from each participant. Unless stated otherwise, all analysis were performed separately for each resting-state run; the identification analysis pooled the four runs of each participant into a single candidate set (Appendix A).

For the ISYB dataset, resting-state fMRI data from healthy Han Chinese adults were analyzed. All participants were right-handed and had no history of neurological or psychiatric disorders. The study was approved by the Ethics Committee of the Institute of Psychology, Chinese Academy of Sciences, and written informed consent was obtained from each participant. Exclusion criteria included regular smoking, pregnancy, MRI contraindications, or any neurological or psychiatric disorders [93]. The publicly released dataset already contains 215 participants (156 female, 59 male) who passed the data providers' quality control; 26 of the 241 scanned participants were excluded and we applied no further exclusion. The participants were between 18 and 30 years of age (mean age 22.6 years, standard deviation 2.67). Each participant underwent a resting-state BOLD scan.

#### A.2 Data Acquisition

Whole-brain echo-planar imaging (EPI) for the HCP was acquired on a customized 3T Siemens Skyra (Connectome Skyra) scanner at Washington University School of Medicine using a 32-channel head coil, with the following parameters: repetition time (TR) = 720 ms, echo time (TE) = 33.1 ms, flip angle = 52°, bandwidth = 2,290 Hz/pixel, in-plane field of view (FOV) = 208 × 180 mm, 72 slices, voxel size = 2.0 mm isotropic, multiband acceleration factor = 8 [23, 95]. Data were acquired over two days. Two runs of resting-state BOLD data were collected per day, totaling about 29 minutes per day (about 58 minutes in total). The two 14.5-minute resting-state BOLD runs acquired on the same day employed opposite phase-encoding directions (right-to-left and left-to-right). Structural images were also acquired with the following parameters: T1-weighted (0.7 mm isotropic resolution, TR = 2,400 ms, TE = 2.14 ms, flip angle = 8°, in-plane FOV = 224 × 224 mm) and T2-weighted (0.7 mm isotropic resolution, TR = 3,200 ms, TE = 565 ms, variable flip angle, in-plane FOV = 224 × 224 mm). Complete parameters and acquisition details for the HCP dataset have been reported elsewhere [96, 97].

The Whole-brain EPI for ISYB was acquired on a 3T GE MR750 scanner at the Magnetic Resonance Research Center of the Institute of Psychology of the Chinese Academy of Sciences, with the following parameters: matrix = 64 × 64, 36 slices, FOV = 220 × 220 $mm^2$, TR = 2,000 ms, TE = 30 ms, flip angle = 90°, voxel size = 3.4 × 3.4 × 4.0 $mm^3$. Structural images were acquired using a T1-weighted sequence with the following parameters: matrix = 256 × 256, 188 slices, FOV = 256 × 256 $mm^2$, TR = 8.16 ms, TE = 3.18 ms, flip angle = 12°, inversion time (TI) = 450 ms, voxel size = 1.0 × 1.0 × 1.0 $mm^3$ [93]. Each participant underwent a resting-state scan (185 volumes; TR = 2,000 ms). Unlike the HCP protocol, ISYB MRI acquisition did not employ multiband acceleration.

#### A.3 Resting-State fMRI Preprocessing

Resting-state fMRI preprocessing for the HCP included the following steps, closely following the pipeline proposed by the HCP consortium [97]: (i) the "minimal preprocessing" approach, including intensity normalization, phase-encoding direction distortion correction, motion correction, and spatial normalization to a standard template (MSMAll, angular penalty version); (ii) Removal of ICA-FIX artifacts [98]; (iii) regression of 24 motion parameters. The final minimally preprocessed BOLD data were registered in the MNI152 space and mapped to the cortical surface of fsLR-32k. The preprocessed data were represented in the CIFTI file format, which combines surface-based data representation for the cortex and volume-based data representation for subcortical gray matter locations (i.e. grayordinates). The CIFTI grayordinate BOLD time series were subsequently used for further analysis. To remove potential signal instability or artifacts at the beginning of the scan, some studies discard the first several time points of each run before using the preprocessed BOLD time series of the HCP standard [97] for subsequent analysis. In the present study, we performed the subsequent analysis separately for each BOLD run in the resting-state of each participant.

The resting-state fMRI data for ISYB were preprocessed using the Connectome Computation System (CCS) [99, 100]. Functional preprocessing included: (i) discarding the first 5 EPI volumes for signal equilibration; (ii) slice timing correction; (iii) motion correction using MCFLIRT; (iv) Normalization of the global mean intensity in 4D; (v) regression of motion artifacts using ICA-AROMA [101]; (vi) removal of linear and quadratic trends. Structural preprocessing included spatial adaptive non-local means denoising, inhomogeneity field correction, brain extraction based on non-local intracranial cavity extraction (NICE), gray–white matter segmentation, and cortical surface reconstruction. The preprocessed data were mapped to the fsLR-32k cortical surface and converted to CIFTI format using the Ciftify toolbox [102].

#### A.4 Full-Band Signal Retention

Before constructing functional brain networks, a critical signal-level consideration is the treatment of the frequency spectrum of the preprocessed BOLD time series. In conventional resting-state fMRI studies, BOLD signals are empirically band-pass filtered into a narrow range (typically 0.01–0.1 Hz) to mitigate scanner drift and physiological noise. However, this empirical truncation lacks theoretical grounding and inevitably discards biologically meaningful neural oscillations outside this range, especially given that non-neuronal artifacts had already been removed via ICA-based denoising (i.e., ICA-FIX and ICA-AROMA) in our preprocessing pipelines. Therefore, rather than applying arbitrary band-pass filtering, the present study retained full-band BOLD signals across the entire detectable frequency spectrum. The theoretical rationale for this choice is grounded in the linear natural logarithm law (N3L) of neural oscillations [103] and the recently proposed three-level integrative model of spontaneous slow oscillations (SSOs) [29, 104]. According to this integrative framework, full-spectrum BOLD fluctuations (spanning slow-1 to slow-6) are not isolated frequency components but are organized into a unified self-organizing hierarchical system comprising three interactive functional levels: the bottom-up detection level (slow-1 to slow-3) for local sensory and perceptual processing, the computation or representation level (slow-4) serving as the modular core for cognitive operations, and the top-down modulation level (slow-5 and slow-6) for global information integration and executive regulation. Crucially, the model emphasizes that the intrinsic functional architecture of the brain emerges from scale-free nested dynamics and cross-frequency coupling across these hierarchical layers, where slow, long-wavelength oscillations provide a global temporal context to modulate faster localized activities [29, 104]. Retaining the full-spectrum BOLD signals thus avoids artificially severing this cross-scale oscillatory orchestration, allowing the constructed functional networks to faithfully capture the complete, multi-level information processing continuum of the human brain.

#### A.5 Node Definition and Time Series Extraction

Following the retention of full-spectrum BOLD signals at the grayordinate level, we mapped these signals onto specific anatomical parcels to extract node-level time series. The cortical nodes were defined using the Schaefer 400 parcellation atlas [105], which delineates 400 functionally homogeneous cortical regions based on functional connectivity in the resting-state. Furthermore, because large-scale cortical switching is closely correlated with and orchestrated by subcortical dynamics [17, 40], we additionally incorporated Tian's 32-subregion subcortical parcellation, derived from functional connectivity gradients [106]. Using this combined parcellation template in standard CIFTI fsLR-32k space, we computed the mean BOLD time series across all grayordinates within each parcel, yielding 432 node-level time series per participant (400 cortical and 32 subcortical nodes).

#### A.6 Time Series Segmentation

To track time-varying modular reconfigurations within the Spatial Connectivity Cycle Flexibility (SCCF) framework while preserving local stationarity, we segmented the continuous node-level time series into consecutive, non-overlapping time windows, with each window $s$ corresponding to a single temporal layer in the multilayer network. Specifically, for the HCP dataset (1,200 time points, TR = 720 ms), each run was partitioned into $T = 20$ windows of 60 time points each (43.2 seconds); for the ISYB dataset (180 time points after discarding the initial 5 volumes, TR = 2,000 ms), each scan was partitioned into $T = 9$ windows of 20 time points each (40.0 seconds). This windowing scheme is in line with established dynamic network flexibility studies [20, 76, 107], ensuring sufficient temporal samples per window for a reliable correlation estimation.

#### A.7 Functional Connectivity Affinity Matrix Construction

Within each time window $s$, rather than relying solely on raw pairwise correlations, we defined network edge weights by computing a functional connectivity affinity matrix based on connectivity fingerprint similarity [21]. Specifically, we first calculated the conventional pairwise Pearson correlation matrix across all 432 nodes within the window $s$, where each node's whole-brain correlation profile represents its functional connectivity fingerprint. Subsequently, the affinity between any pair of nodes was quantified by computing the Pearson correlation between their respective connectivity fingerprints, which produced a symmetric $432 \times 432$ affinity matrix $A_s$ for each temporal layer.

This two-step approach is grounded in connectome generative models and the principle of connectivity homophily [26, 27, 108], which posit that similarity in whole-brain connectivity patterns reflects the functional proximity of two regions more faithfully than direct pairwise coupling alone. By integrating temporal synchrony with spatial topological homophily into a single metric, the resulting affinity matrix $A_s$ naturally encompasses positive and negative weights across a continuous spectrum of strengths.

#### A.8 Retention of Signed and Weak Connections

To construct the multilayer network from the window-specific affinity matrices $\{A_s\}$, we deliberately preserved the complete, signed, and fully connected structure of each matrix, retaining both positive and negative connections without applying any sparsification threshold.

The retention of signed (both positive and negative) weights is anchored in a central hypothesis of the present study: whole-brain dynamic modular reconfigurations are fundamentally driven by the ongoing interplay between cooperative (positive) and antagonistic (negative) interactions. Intrinsically, macroscale brain systems exhibit competitive dynamics, most prominently exemplified by suppression and antagonism between the default mode network (DMN) and the action mode network (AMN) [9, 24, 25], along with dynamic calibration between antagonism and cooperation across transmodal control systems [36, 37]. At the neurobiological level, negative functional weights capture meaningful functional inhibition or phase delays among neuronal populations [30, 109], providing the competitive push–pull forces that propel boundary and relay regions to switch modular allegiances over time. Restricting networks to exclusively positive connections would obscure these antagonistic drivers of network dynamics. Importantly, because we refrained from global signal regression (GSR) during preprocessing, a step known to mathematically induce spurious anticorrelations—the negative affinities in $A_s$ reflect genuine physiological antagonism, enabling their rigorous incorporation into signed multilayer modularity maximization [19, 22, 30].

Similarly, we refrained from imposing any sparsification threshold on $A_s$, thus preserving weak connections for two key reasons. First, from a methodological point of view, selecting an arbitrary sparsity threshold (e.g., retaining only the top 10% of edges) lacks a consensus criterion and can substantially distort network topology, compromising cross-study comparability and reproducibility. Second, and more critically from a neurobiological point of view, weak connections below conventional thresholds act as vital topological bridges that sustain long-range information integration and support higher-order cognition [110]. Truncating these weak affinities would artificially fragment the network and sever subtle cross-tier interactions. Consequently, characterizing each temporal layer by its intact signed affinity matrix $A_s$ circumvents arbitrary thresholding biases while preserving the richest topological foundation for tracking dynamic community reconfigurations.

#### A.9 Multilayer Network Community Detection

To track dynamic community structure while preserving both synergistic and antagonistic interactions, we represented each time window as a signed network layer and organized the sequence of temporal layers into a multilayer network within the generalized modularity maximization framework proposed by Mucha et al. [19]. Specifically, the dynamic functional connectivity affinity matrix $A_s$ from the $s$-th time window constitutes the intralayer structure of the $s$-th temporal layer, where positive and negative affinities form signed intralayer edges (Fig. A1c). Across adjacent temporal layers, each brain region is linked to itself via interlayer coupling edges to capture the temporal continuity of community assignments across consecutive time windows. The multilayer community detection was then performed using the generalized Louvain algorithm (GenLouvain) [19] (Fig. A1d).

Building upon this multilayer formulation, we further adopted the community stability framework based on a continuous-time Markov process proposed by Lambiotte et al. [83] to evaluate the stability of community structures across scales. In this framework, each brain region within a given temporal layer defines a network state; a continuous-time random walker can traverse between brain regions within the same layer via intralayer edges or transition between adjacent temporal layers via interlayer coupling. The stability of a given community partition is quantified by examining the intralayer retention characteristics of the random walk across different Markov time scales. To faithfully account for the signed nature of our affinity networks, we incorporated both positive synergistic edges and negative antagonistic edges into the community stability evaluation, ensuring that the resulting community partitions are simultaneously shaped by cooperative attraction within modules and competitive repulsion between modules.

In summary, by integrating signed multilayer modularity maximization [19] with the continuous-time Markov stability framework [83], this approach simultaneously

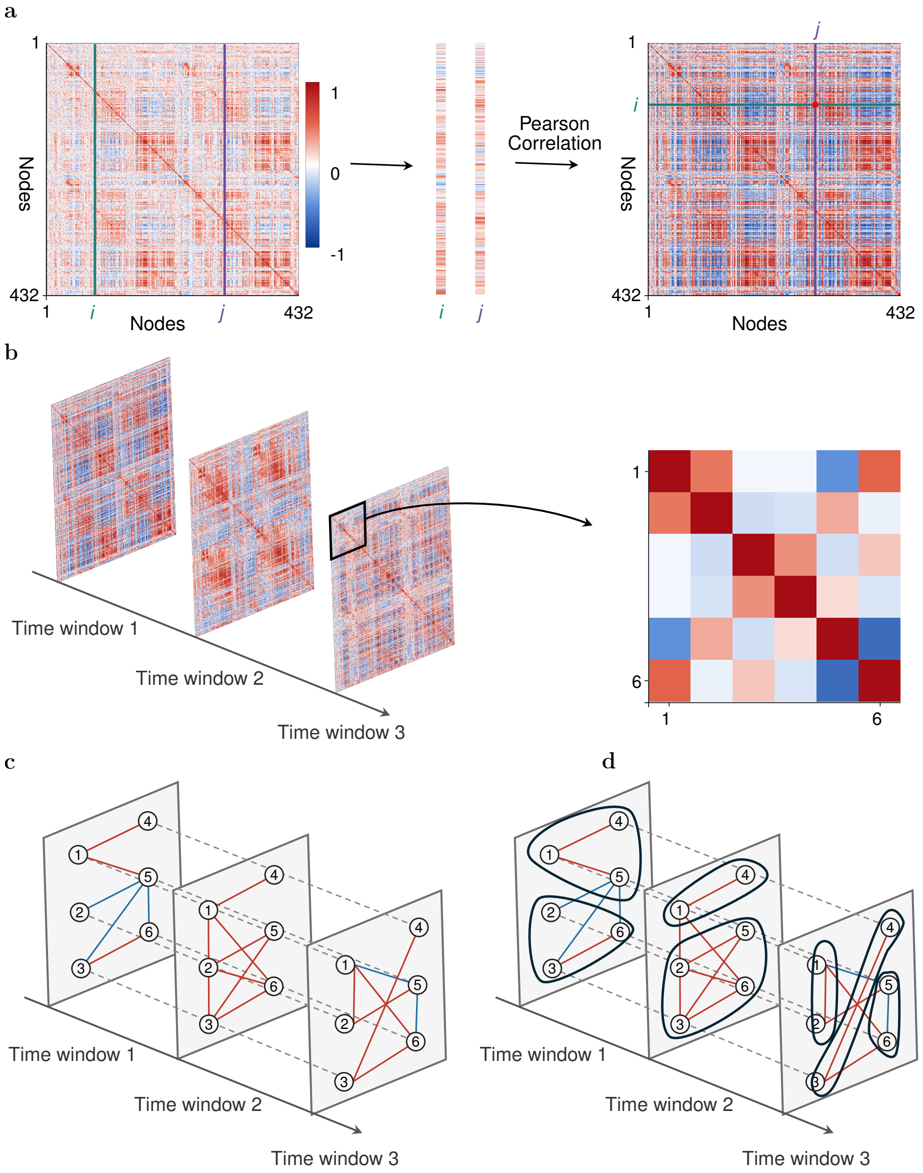


**Fig. A1** Construction of dynamic functional connectivity affinity networks and multilayer community detection. **a**, The connectivity fingerprints of nodes $i$ and $j$ in the DFC matrix are correlated using Pearson correlation, giving the $(i, j)$ element of the affinity matrix (red dot). **b**, Affinity matrices from successive time windows form the temporal layers of a multilayer network (three of the $T$ time windows are shown; $T = 20$ for HCP and $T = 9$ for ISYB); the $6 \times 6$ submatrix of time window 3 is enlarged. **c**, The signed networks of the three time windows shown in **b**, with positive and negative affinities shown as positive and negative intralayer edges (red and blue); dashed grey edges denote the interlayer coupling of each region to itself across adjacent windows. **d**, Multilayer community detection using the GenLouvain algorithm, incorporating interlayer coupling between corresponding nodes across adjacent layers; outlines denote the detected communities.

characterizes intralayer functional synergy and antagonism, interlayer community continuity across consecutive time windows, and the multiscale stability of dynamic community structures within a unified network model.

#### A.10 Markov Process in Multilayer Networks

Building upon the aforementioned concept of community stability, we formulated the random walk in the multilayer network as a continuous-time Markov process and employed Markov stability to quantify the stability of a given community partition across different dynamical time scales. Specifically, community stability is defined as the deviation of the intralayer retention of random walks at a finite time scale from the long-term random expectation:

$$R(t) = \sum_{C} [P(C,t) - P(C,\infty)] \tag{A1}$$

where $t$ denotes the time scale of the random walk, $P(C,t)$ represents the joint probability that a random walker is located within community $C$ at both the initial state and after time $t$, and $P(C,\infty)$ denotes the long-term random expectation of the same event when the random walk reaches the stationary state. Therefore, $R(t)$ reflects the extent to which the random walk maintains its community membership under a given community partition at a finite time scale. A larger $R(t)$ indicates that the random walker tends to remain within the original community at that time scale, signifying a higher stability of the community partition at the corresponding time scale.

For a multilayer network comprising $N$ nodes and $T$ time layers, the node $i$ in layer $s$ is represented as a composite state $(i,s)$. Since the functional connectivity affinity simultaneously contains positive and negative connections, we first decomposed the signed affinity matrix $A_s$ of each time layer into two non-negative components:

positive synergistic connections and negative antagonistic connections, such that its element $A_{ijs} = A^+_{ijs} - A^-_{ijs}$ (self-connections are excluded, $A_{iis} = 0$). Here, $A^+_{ijs}$ denotes the positive connection strength between nodes i and j in layer s, reflecting the synergistic association between brain regions, whereas $A^-_{ijs}$ denotes the absolute strength of negative connections, reflecting the antagonistic relationship between brain regions.

Within this state space, a random walker can move between different brain regions along the functional connections within the same time layer, or transition from one time layer to the same brain region in another time layer via interlayer coupling. Considering that positive connections provide synergistic flow and negative connections impose antagonistic constraints, the transition weight from state $(j, r)$ to state $(i, s)$ is expressed as $(A^+_{ijs} - A^-_{ijs})\delta_{sr} - \delta_{ij}C_{jsr}$. Specifically, $A^+_{ijs}\delta_{sr}$ represents the positive synergistic connection within the same time layer, while $A^-_{ijs}\delta_{sr}$ represents the absolute strength of the negative antagonistic connection. A random walker can only move from node $j$ to node $i$ through the functional connection in layer s when the two nodes are in the same layer (i.e., $s = r$). Meanwhile, $-\delta_{ij}C_{jsr}$ represents the interlayer coupling, which links only the same node across different layers (i.e., $i = j$).

For the multilayer state $(j, r)$, the positive and negative connection strengths within its r-th time layer are defined as $k^+_{jr} = \sum_i A^+_{ijr}$ and $k^-_{jr} = \sum_i A^-_{ijr}$, respectively. The intralayer connection strength is thus defined as $k_{jr} = k^+_{jr} - k^-_{jr}$. Meanwhile, the interlayer coupling strength between state $(j, r)$ and other time layers is defined as $c_{jr} = \sum_s C_{jsr}$. The total transition weight out of state $(j, r)$ is then $\kappa_{jr} = k_{jr} - c_{jr}$.

To describe the continuous evolution of the random walk over time, let $p_{is}(t)$ denote the probability that the random walker is in state $(i, s)$ at time $t$. For a continuous-time Markov process, the rate of change of the probability of a given state is jointly determined by the probability inflow from other states into that state and the probability outflow from that state to others. Therefore, the rate of change for state $(i, s)$

can be expressed as:

$$\dot{p}_{is} = \sum_{jr} \left( \frac{(A^{+}_{ijs} - A^{-}_{ijs})\delta_{sr} - \delta_{ij} C_{jsr}}{\kappa_{jr}} \right) p_{jr} - p_{is} \tag{A2}$$

Defining the Laplacian operator of the multilayer network as

$$L_{ijsr} = \frac{(A^{+}_{ijs} - A^{-}_{ijs})\delta_{sr} - \delta_{ij} C_{jsr}}{\kappa_{jr}} - \delta_{ij}\delta_{sr} \tag{A3}$$

the aforementioned differential equation can be written in matrix form as $\frac{d\boldsymbol{p}}{dt} = \mathbf{L}\boldsymbol{p}$. The corresponding stationary state can be expressed as $p^{*}_{jr} = \frac{\kappa_{jr}}{2\mu}$, where $2\mu = \sum_{jr} \kappa_{jr}$ represents the total transition weight of the multilayer states. Applying a first-order Taylor expansion approximation to the solution of the differential equation, $\boldsymbol{p}(t) = e^{t\mathbf{L}}\boldsymbol{p}(0) \approx (\mathbf{I} + t\mathbf{L})\boldsymbol{p}(0)$, yields the actual conditional transition probability from the starting state $(j, r)$ to the ending state $(i, s)$ as $p_{is|jr} = \delta_{ij}\delta_{sr} + tL_{ijsr}$.

Given a community partition $g_{is}$, assume the random walker is initially located at state $(j, r)$. The transition probability of being at state $(i, s)$ after time $t$ is then $p_{is|jr}$. When the two states belong to the same community, $\delta(g_{is}, g_{jr}) = 1$; otherwise, it is 0. Therefore, summing over all possible initial and final states yields the joint probability that the random walker remains within its initial community after time $t$ as:

$$P(C, t) = \sum_{ijsr} (\delta_{ij}\delta_{sr} + tL_{ijsr}) p^{*}_{jr} \delta(g_{is}, g_{jr}) \tag{A4}$$

As $t \to \infty$, the random walk reaches the stationary state. At this point, the expected joint probability of the random walk from state $(j, r)$ to state $(i, s)$ can be expressed as:

$$P(C, \infty) = \sum_{ijsr} \rho_{is|jr} p^{*}_{jr} \delta(g_{is}, g_{jr}) \tag{A5}$$

where the probability of the walker being at state $(j, r)$ in the stationary state is $p^*_{jr} = \frac{\kappa_{jr}}{2\mu}$.

To characterize the expectation of the random walk as $t \to \infty$, we decompose the walk starting from state $(j, r)$ into three parts based on the connection types in the multilayer signed network: intralayer positive connections, intralayer negative connections, and interlayer coupling, and define their expected contributions accordingly. For state $(j, r)$, the signed shares of the walker moving along intralayer positive connections, intralayer negative connections, and interlayer coupling are $\frac{k^+_{jr}}{\kappa_{jr}}, -\frac{k^-_{jr}}{\kappa_{jr}}$, and $-\frac{c_{jr}}{\kappa_{jr}}$, respectively. For the intralayer positive part, within the same time layer $r = s$, the expected probability of the walker reaching state $(i, s)$ is $\frac{k^+_{is}}{2m^+_s}\delta_{sr}$, where $2m^+_s = \sum_{ij} A^+_{ijs}$ denotes the total positive weight of layer s. Similarly, the expected probability corresponding to intralayer negative connections is $\frac{k^-_{is}}{2m^-_s}\delta_{sr}$, where $2m^-_s = \sum_{ij} A^-_{ijs}$ denotes the total negative weight of layer s. For the interlayer part, the random walker can only transition between different time layers via $C_{jsr}$, and this transition must occur between the same brain region, thus requiring $i = j$. The corresponding interlayer expected contribution is represented by $-\delta_{ij}\frac{C_{jsr}}{\kappa_{jr}}$.

Therefore, the conditional transition probability $\rho_{is|jr}$ can be expressed as:

$$\rho_{is|jr} = \left( \frac{k^+_{is}}{2m^+_s} \frac{k^+_{jr}}{\kappa_{jr}} - \frac{k^-_{is}}{2m^-_s} \frac{k^-_{jr}}{\kappa_{jr}} \right) \delta_{sr} - \delta_{ij} \frac{C_{jsr}}{\kappa_{jr}} \tag{A6}$$

Substituting the above components into $R(t)$, the community stability can be further decomposed into three parts: intralayer positive/negative connections, their corresponding random expectations, and interlayer coupling:

$$\begin{aligned} R(t) = \frac{1}{2\mu} \sum_{ijsr} \Bigg\{ & \left[ t(A^+_{ijs} - A^-_{ijs}) - \left( \frac{k^+_{is}k^+_{jr}}{2m^+_s} - \frac{k^-_{is}k^-_{jr}}{2m^-_s} \right) \right] \delta_{sr} \\ & + (1-t)\delta_{ij}C_{jsr} + (1-t)\delta_{ij}\delta_{sr}\kappa_{jr} \Bigg\} \delta(g_{is}, g_{jr}) \end{aligned} \tag{A7}$$

Among these, the last term $(1-t)\delta_{ij}\delta_{sr}\kappa_{jr}$ is non-zero only when $i=j$ and $s=r$. In this case, the two states belong to the same community; thus, summing this term over all states yields only a constant $1-t$ that is independent of the community partition and does not affect the relative stability between different community partitions. Therefore, in the process of maximizing community stability, this constant term independent of the community partition can be ignored, leading to the multilayer signed modularity function for community partition optimization:

$$
\begin{aligned}
Q(t) = \frac{1}{2\mu}\sum_{ijsr}\Bigg\{ & \left[t(A^{+}_{ijs} - A^{-}_{ijs}) - \left(\frac{k^{+}_{is}k^{+}_{jr}}{2m^{+}_{s}} - \frac{k^{-}_{is}k^{-}_{jr}}{2m^{-}_{s}}\right)\right]\delta_{sr} \\
& + (1-t)\delta_{ij}C_{jsr}\Bigg\}\delta(g_{is}, g_{jr})
\end{aligned} \tag{A8}
$$

Thus, the intralayer term in $Q(t)$ simultaneously incorporates the difference between the actual signed connections and their corresponding random expectations, where positive connections provide a positive contribution to the synergistic associations within communities, while negative connections impose a negative contribution to the co-membership of brain regions with antagonistic relationships. The interlayer coupling term reflects the continuity of community membership for the same brain region across adjacent time layers via $(1-t)C_{jsr}$.

For a given time $t$, community stability measures the degree to which the random walk remains within the same community at a given Markov time scale, relative to the random walk expectation as $t \to \infty$. Therefore, maximizing $Q(t)$ is equivalent to finding a community partition that makes the random walk more inclined to stay within communities. For the multilayer signed network constructed in this study, this implies that the community partition obtained by maximizing $Q(t)$ is both consistent with the signed functional connection structure of each time layer and capable of maintaining the continuity of community membership across adjacent time layers.

#### A.11 Configuration of Interlayer Connections

Following the derivation of the multilayer signed modularity function, we further determined the specific values for the interlayer connection matrix $C$ and the Markov time parameter $t$. $C_{jsr}$ denotes the interlayer coupling of node $j$ between time layers $s$ and $r$, which controls the community continuity of the same brain region across different time layers. We set $C_{jsr} = 1$ when the two layers are adjacent ($|s-r| = 1$) and $C_{jsr} = 0$ otherwise, for every node $j$, so that $C$ is symmetric ($C_{jsr} = C_{jrs}$). $C_{jsr}$ is therefore a coupling magnitude whose sign is supplied by the derivation, entering the transition weights as $-\delta_{ij}C_{jsr}$ and $Q(t)$ as $+(1-t)\delta_{ij}C_{jsr}$. Unit coupling is the conventional choice; because the affinity weights are bounded by unity, the resulting interlayer term $(1-t)C_{jsr}$ remains on a comparable scale with a single intralayer weight. With this choice, $c_{jr} = \sum_s C_{jsr}$, and hence $\kappa_{jr} = k_{jr} - c_{jr}$, equals 2 for interior layers and 1 for the first and last layers. Restricting the coupling to adjacent layers ensures that it reflects local temporal continuity rather than long-range temporal association.

This study further set the Markov time parameter $t = 1/1.1$. Under this setting, $(1-t)C_{jsr} > 0$, so maintaining the same community membership for the same brain region in adjacent time layers yields a positive contribution, thereby reflecting a certain degree of temporal continuity in the dynamic community partition. Dividing $Q(t)$ by $t$, which leaves the maximizing partition unchanged, writes the same objective in the standard multilayer form, with the intralayer null model scaled by the resolution parameter $\gamma$ and the interlayer term scaled by the coupling parameter $\omega$:

$$\frac{Q(t)}{t} = \frac{1}{2\mu}\sum_{ijsr}\left\{\left[(A^{+}_{ijs} - A^{-}_{ijs}) - \gamma\left(\frac{k^{+}_{is}k^{+}_{jr}}{2m^{+}_{s}} - \frac{k^{-}_{is}k^{-}_{jr}}{2m^{-}_{s}}\right)\right]\delta_{sr} + \omega\,\delta_{ij}C_{jsr}\right\}\delta(g_{is}, g_{jr}) \tag{A9}$$

where $\gamma = 1/t = 1.1$ and $\omega = (1-t)/t = 0.1$ are the conventional resolution and interlayer coupling values used in multilayer modularity studies of brain functional networks [20, 111, 112].

#### A.12 GenLouvain Community Optimization

Following the construction of the multilayer modularity function and the intralayer and interlayer connections described above, we used the GenLouvain algorithm to maximize the multilayer modularity function $Q(t)$ for each multilayer network—that is, for each resting-state run of each participant. The algorithm was implemented using the GenLouvain package for MATLAB (https://github.com/GenLouvain/GenLouvain). Modularity optimization assigns brain regions to communities across time layers, yielding the community affiliation of each brain region at each time layer (Fig. A1d).

Because modularity optimization is stochastic, different optimization runs on the same multilayer network may yield different community partitions. To reduce the dependence of the resulting partition on any single optimization run, we independently performed 100 GenLouvain optimizations for each multilayer network, generating 100 candidate community partitions. We then quantified the agreement between candidate partitions with the $z$-scored Rand coefficient, a chance-corrected measure of partition similarity, and selected a representative partition. Specifically, we first calculated the pairwise $z$-scored Rand similarity between all 100 candidate partitions. For each candidate partition, we then calculated its mean $z$-Rand similarity with the other 99 candidate partitions and selected the partition with the highest mean $z$-Rand similarity as the representative community partition for the corresponding multilayer network. The representative partitions were used for all subsequent analyses, including node flexibility and the community co-occurrence matrix from which the DMCF is derived. This procedure was implemented using the Consensus and Comparison Methods provided in the MATLAB Network Community Toolbox (http://commdetect.weebly.com).

Following these procedures, each resting-state run of each participant was summarized by a community assignment for every node in every time window.

#### A.13 Flexibility Atlas

Based on the representative community assignments obtained above, a node may change its community affiliation across consecutive time layers of the multilayer network. The frequency of such community switching over time characterizes a region's dynamic reconfiguration and is defined as node flexibility [20]:

$$\text{Node flexibility} = \frac{\text{Number of community switches}}{T-1} \tag{A10}$$

where $T$ denotes the total number of time layers and $T-1$ represents the total number of possible transitions between consecutive layers. Higher node flexibility indicates more frequent participation in dynamic community reorganization.

According to the mode of community switching, node flexibility can be further decomposed into disjoint flexibility and cohesive flexibility [113]. Disjoint flexibility quantifies the frequency with which a node switches to another community on its own, that is, no other node in the same layer makes the same community-to-community transition, whereas cohesive flexibility quantifies the frequency with which the node's transition is shared by at least one other node, thereby capturing local independent adjustments and coordinated multi-node reorganization, respectively. All flexibility measures were computed for each resting-state run using the Dynamic Community Structure module in the MATLAB Network Community Toolbox (http://commdetect.weebly.com). Individual node flexibility maps were then averaged across runs within each participant and subsequently across participants to construct the group-level flexibility atlas. The resulting group-level whole-brain flexibility atlas is presented in Appendix B, Section B.1 (Fig. B1).

#### A.14 Community Co-occurrence Matrix

Node-level flexibility captures how often each region changes its community affiliation, but it reduces each region to a single number and discards the pairwise structure that describes which regions are grouped together at any moment. To preserve this structure, we summarized the representative community assignments of each participant-run as a community co-occurrence matrix, which records, for every pair of regions, the proportion of time layers in which the pair belongs to the same functional module.

For nodes $i$ and $j$, the co-occurrence frequency $P_{ij}$ is the proportion of time layers in which the two nodes are assigned to the same community:

$$P_{ij} = \frac{1}{T} \sum_{t=1}^{T} \delta(g_{it}, g_{jt}) \tag{A11}$$

where $g_{it}$ and $g_{jt}$ denote the communities to which nodes $i$ and $j$ belong in layer $t$, $T$ is the number of time layers in the run (20 for each HCP run and 9 for the ISYB run), and $\delta(g_{it}, g_{jt})$ equals 1 when the two nodes share a community in that layer and 0 otherwise.

Repeating this calculation for all node pairs yields a symmetric $432 \times 432$ matrix ($P_{ij} = P_{ji}$) whose entries lie in $[0, 1]$; the diagonal equals 1 by construction, because a node shares a community with itself in every layer. For each participant, the matrices of the available runs were averaged, and these participant-level matrices were then averaged with equal weight per participant, giving a group-level co-occurrence matrix in which each entry is the probability that two regions belong to the same functional module. This group-level matrix is shown as a heat map in Fig. 1d.

#### A.15 Group-level Co-occurrence Network

To visualize how functional systems were linked by their strongest community co-occurrences, we represented the group-level co-occurrence matrix (Section A.14) as a sparse network (Fig. 1e). Specifically, the 93,096 upper-triangular entries (unique node pairs) of the $432 \times 432$ group-level matrix were ranked by co-occurrence probability, and added in descending order until every Du-15 functional network had at least one connection to another network; the selected edges formed a sparse adjacency matrix.

For visualization, the 432 nodes were arranged on a circle using MNE-Python and MNE-Connectivity [114], such that each functional system occupied a contiguous arc. Nodes were ordered around the ring by their Du-15 functional network, followed by the 32 Tian subcortical parcels and the 28 unassigned cortical parcels. Nodes were coloured by their functional group, and the retained edges were coloured according to their co-occurrence probability on a viridis scale spanning the minimum and maximum values among the selected edges.

#### A.16 Maximum-Overlap Community Map

To bridge the gap between dynamic community organization and canonical static networks, we distilled the group-level co-occurrence matrix (Section A.14) into a discrete consensus partition, termed the maximum-overlap community map (Fig. 1g). Because this group-level overlap probability directly reflects how frequently different brain regions integrate with one another across time windows and participants, we applied spectral clustering to the matrix to identify communities of regions that consistently group together across cortical and subcortical structures. We explored this network hierarchy across multiple spatial scales: a coarse two-system division, an intermediate three-community partition that captures an emerging transitional zone, and a finer seven-community partition directly comparable to the canonical Yeo reference networks. For visualization, the cortical results were projected onto the brain surface alongside the reference atlas (Fig. 1g). Ultimately, this map serves as a static, low-dimensional summary of the dynamic co-occurrence landscape.

To evaluate the alignment between our finest partition and the canonical reference, we matched each non-limbic reference network to the consensus communities that encompassed the majority of its cortical parcels (breaking ties in favor of smaller communities). Excluding the limbic regions, the overall overlap rate was calculated as the proportion of successfully matched parcels across the cortex. Notably, because a single consensus community could correspond to multiple reference networks, this metric reflects overall spatial coverage rather than a strict one-to-one correspondence.

#### A.17 Subcortical Co-Occurrence Pattern Maps

To characterize how individual subcortical structures dynamically couple with the cerebral cortex, we extracted the subcortical routing profiles directly from the group-level community co-occurrence matrix (Section A.14). Because this matrix captures the pairwise co-occurrence probability across all brain regions, the profile associated with a given subcortical nucleus directly reflects its whole-brain integration preference. We visualized each subcortical co-occurrence pattern using two complementary views (Fig. 2c–e; Fig. B5): first, its cortical co-occurrence probabilities were projected onto the brain surface to map its spatial topography across the cortex; second, its strongest couplings were rendered as a circular connectogram, grouped by canonical functional networks, to highlight its primary integration partners. Because these profiles are derived directly from the group-level co-occurrence matrix rather than estimated at the individual level, the resulting maps provide a direct readout of the conserved group-level co-occurrence architecture.

#### A.18 Dynamic Modular Co-occurrence Fingerprint

Because the matrix is symmetric and its diagonal is constant, only the off-diagonal entries carry meaningful information. As in functional connectivity fingerprinting [21], we retained the strict upper triangle (excluding the diagonal) in the row to form the DMCF vector.

$$\mathrm{DMCF} = (P_{12}, \ldots, P_{1n}, P_{23}, \ldots, P_{2n}, \ldots, P_{(n-1)n}) \tag{A12}$$

For $n = 432$ nodes, this vector comprises $n(n-1)/2 = 93{,}096$ entries, each representing the frequency of co-occurrence of a given pair of nodes to provide an individual-specific description of modular organization.

Using the DMCF defined above (Fig. A2), we tested whether the fingerprint of a single run carries a stable individual signature. For the 917 participants who completed all four resting-state runs, each run was treated as a separate fingerprint (3,668 fingerprints in total), and pairwise Pearson correlations were computed across all 93,096 co-occurrence values. Excluding the query run itself from the candidate pool, each run was assigned to the participant yielding the highest correlation. The identification was scored as correct when this rank-1 nearest neighbor belonged to the same participant. This nearest-neighbor assignment was accurate for 2,292 of the 3,668 runs (62.49%), far exceeding the analytic chance level of 0.08% (3 out of 3,667, reflecting the three remaining runs of the same participant). Thus, the DMCF carries a robust and reproducible individual signature.

DMCF was subsequently used as the characteristic vector for the analysis of behavioral associations (Fig. A3).

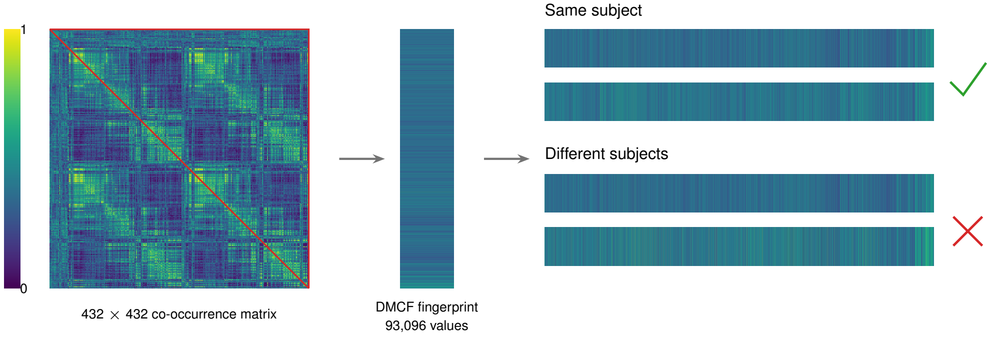


**Fig. A2 Cross-run identification of individual participants using the DMCF.** Left, the 432 × 432 community co-occurrence matrix of one representative resting-state run (colour bar, co-occurrence frequency); the red outline marks the strict upper triangle that is retained, because the matrix is symmetric and its diagonal is constant. Middle, these 93,096 co-occurrence values, flattened in row order, form the DMCF fingerprint of that run. Right, each of the four resting-state runs of the 917 HCP participants who completed all four runs was treated as a separate fingerprint (3,668 in total), and its Pearson correlation with every other fingerprint was computed. The query run is shown against its rank-1 nearest neighbour, which belongs to the same participant (tick), and against the most similar run of a different participant (cross); each run was assigned to the participant with the highest correlation among all other runs, with the run itself excluded. This assignment was correct for 2,292 of 3,668 runs (62.49%), against an analytic chance level of 0.08% (3/3,667).

#### A.19 Behavioral target selection and factor analysis

Rather than specifying target behaviors a priori, we identified candidate measures through an initial data-driven screening that combined preliminary association analyzes with machine learning. Specifically, we applied exploratory 10-fold cross-validated connectome-based predictive modeling (CPM) to the features of DMCF (after regressing age and sex within each training fold) and retained behaviors whose pooled out-of-fold predictions were significantly correlated with the observed values ($p < 0.05$, uncorrected). To ensure a less redundant and more independent behavioral space prior to modeling, we also incorporated manual curation alongside statistical criteria such as requiring alignment with the primary axis of the behavioral battery for a subset of measures and collapsing six highly collinear delay-discounting variants into a single measure. This hybrid screening procedure yielded 29 candidate behaviors that showed preliminary predictability of the DMCF. (Note that because this exploratory selection was performed within the same cohort, the resulting targets serve to anchor our characterization of brain–behavior relationships rather than to provide an unbiased benchmark of out-of-sample generalization.) The screening counts and the composition of each factor are reported in Appendix B.5.

To further summarize these 29 candidate measures and account for their shared variance, we submitted them to an exploratory factor analysis (EFA). Before EFA, all measures were residualized for age and sex, imputed for minor missing data ($k$-nearest-neighbor imputation, $k = 10$, after excluding participants missing $> 50\%$ of the measures; final $N = 868$), and $z$-standardized. Using maximum likelihood extraction with varimax rotation, we identified eight latent factors based on the Kaiser criterion (eigenvalues $> 1$). In particular, eight of the 29 behaviors exhibited low loadings in all extracted factors (maximal $|\text{loading}| < 0.30$) and were therefore retained as distinct independent components. Together, the eight latent factors and eight independent components parsimoniously decomposed the behavioral space into 16 target dimensions for all subsequent analysis.

#### A.20 Connectome-based behavioral prediction

To evaluate whether the whole-brain DMCF profile predicts individual differences across the 16 behavioral targets, we employed linear kernel ridge regression (KRR) using all 93,096 upper-triangular co-occurrence features without prior edge selection. The similarity between the participants was operationalized through the linear kernel $K = XX^\top$. Prediction performance was evaluated using a nested cross-validation framework consisting of an outer 10-fold loop for generalization assessment and an inner 5-fold loop for hyperparameter tuning. To prevent data leakage, age and sex were regressed out within each outer fold: nuisance regression parameters were estimated strictly on the training set and applied to residualize both the training and held-out test targets, as well as the corresponding blocks of the kernel matrix. Within each inner loop, the $L_2$ regularization parameter $\lambda$ was optimized on a 17-point logarithmic grid ($\log_{10} g \in [-6, 2]$ in steps of 0.5, scaled by $\mathrm{tr}(K)/n_{\mathrm{train}}$) to maximize the pooled inner-fold Pearson correlation.

The Out-of-fold prediction accuracy ($R_{\mathrm{OOF}}$) was quantified as the Pearson correlation between the pooled holdout predictions across all 10 outer folds and the residualized observed scores (using complete cases per target: $N = 828$ for latent factors; $N = 991$–1,031 for independent components). Statistical significance was established via non-parametric permutation testing (1,000 permutations), in which target values were randomly shuffled across participants and the entire nested KRR pipeline was re-executed. The One-sided permutation values $p$ ($p_{\mathrm{perm}}$) were corrected for multiple comparisons using the Benjamini–Hochberg false discovery rate (FDR) procedure ($q_{\mathrm{BH}} < 0.05$) separately within the eight latent factors and the eight independent components. To verify robustness against scan duration, the entire prediction pipeline was repeated in a sensitivity cohort restricted to participants with all four resting-state runs ($N = 779$ for latent factors; $N = 908$–917 for independent components; 200 permutations).

#### A.21 Behavioral association analysis

To characterize how inter-regional integration relates to individual behavioral differences, we performed an edge-level association analysis across behavioral targets (including the 29 candidate measures, the eight latent factors, and general intelligence $g$). For each target, all 93,096 upper-triangular DMCF edges and the behavioral score were residualized for age and sex using ordinary least squares. We then computed the Pearson correlation between each edge and the target, retaining the 500 most positively correlated and 500 most negatively correlated edges as disjoint sets to capture the inter-network integration frequency features most strongly associated with each behavioral measure.

General intelligence $g$ was estimated after the approach of Dubois et al.[62]: ten HCP cognitive measures (picture vocabulary, oral reading recognition, picture sequence memory, dimensional change card sort, pattern comparison processing speed, Penn progressive matrices, variable short Penn line orientation, Penn word memory, flanker inhibitory control and attention, and list sorting working memory) were $z$-scored and submitted to a hierarchical exploratory factor analysis with maximum-likelihood extraction and four correlated group factors (oblimin rotation); the general factor of the resulting Schmid–Leiman solution was scored by regression, and each participant's score on this factor was used as $g$ throughout, including for the whole-brain connectograms of Section B.6. All ten measures were positively loaded on the general factor, so higher scores indicate better performance. The score is centred but not unit-variance ($SD = 0.89$) and was not residualized for age or sex at construction; as for every other target, residualization was performed at analysis time using exact age and sex. The model was estimated in 875 participants with complete cognitive data; 874 had a usable participant identifier, 834 of whom also had resting-state imaging, and 785 all four runs.

Because each DMCF edge quantifies the frequency with which two brain nodes are integrated into the same functional community, these features are inherently additive. We therefore subtracted the summed frequencies of the negative edge set from those of the positive edge set (Total = Pos − Neg) to derive an individual-level summary score of net integration frequency (Fig. A3). The composite frequency score and the behavioral measure were then residualized for age and sex and correlated through Pearson $R$, reflecting how the overall dynamic integration between the distributed regions of the brain collectively gives rise to a given behavior.

For behaviors showing clear associations, we further examined which specific functional networks harbored these behaviorally relevant integration features by projecting the top 1,000 edges (500 positive, 500 negative) onto whole-brain circular connectograms for subsequent network-level analysis.

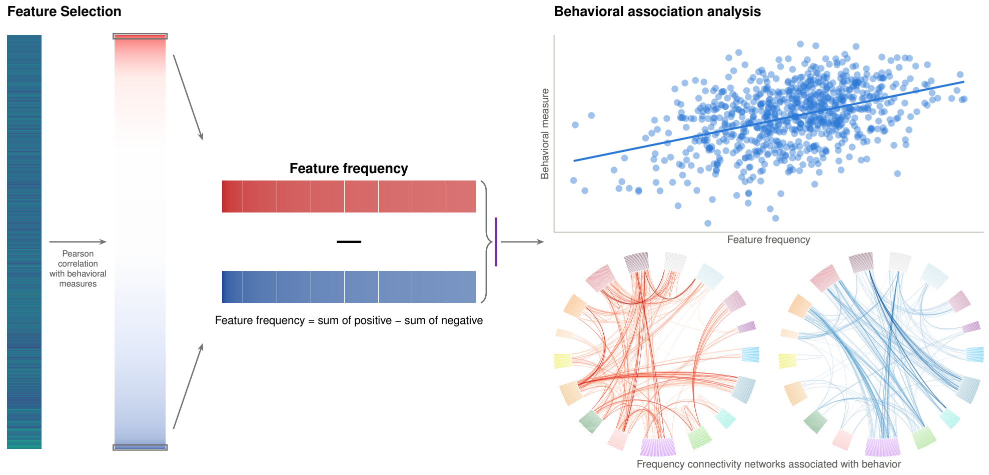


**Fig. A3 Feature selection and behavioral association of the DMCF. Left**, the 93,096 upper-triangular DMCF edges of the 432-node atlas, each residualized for age and sex, are correlated with the residualized behavioral measure and sorted in descending order (top, $r = +0.182$; bottom, $r = -0.207$). The 500 most positive and the 500 most negative edges (boxed) are expanded into two 500-cell strips and combined into a one-dimensional feature frequency (sum of positive minus sum of negative). **Right**, for the 785 HCP participants with all four resting-state runs, the residualized feature frequency is plotted against the residualized $g$ score (Pearson $r = 0.424$, $R^2 = 0.180$, $p = 1.27 \times 10^{-35}$; line, ordinary least-squares fit), and the 500 positive and 500 negative edges are shown as whole-brain circular connectograms (red, positive; blue, negative). Nodes follow a fixed 15-group order (DN-A, DN-B, LANG, AUD, FPN-B, FPN-A, dATN-A, SAL/PMN, AMN, dATN-B, PM-PPr, SMOT-A, SMOT-B, VIS-C, VIS-P, subcortical and uncertain), comprising 400 Schaefer 2018 cortical parcels and 32 Tian subcortical nuclei; node colour indicates network membership (cortical colours from the Zuo LGP atlas, light blue for subcortical nuclei, light grey for nodes outside the 15-network Du-15 scheme), and edge colour encodes $|r|$ on each panel's own scale, so colour depth is not comparable between the two panels. Edge selection and correlation use the same participants, so all values shown are in-sample effect sizes rather than cross-validated prediction accuracy. Correlations were computed with $g$ as an example behavioral measure; the same procedure applies to any behavioral measure.

#### A.22 Subcortical participation and key-node association analysis

To characterize the subcortical contributions to general intelligence ($g$) and the three retained personality dimensions (agreeableness, extraversion, and openness; Section B.7), we first constructed subcortical participation maps. For each behavioral target, all 93,096 upper-triangular DMCF edges were correlated with the behavioral score after residualizing both variables for age and sex. To preserve distributed node–network coupling patterns without omitting informative connections, edges passing a feature-selection threshold of $p < 0.05$ were retained and partitioned into positive and negative sets according to the sign of their correlation. For each of the 32 Tian subcortical nuclei, we then quantified nodal participation (degree) by counting the number of retained positive and negative edges incident to that nucleus, summed across the left and right hemispheres.

Next, for each target, we identified the subcortical nuclei with the highest positive and negative degrees as key seed nodes and mapped the cortical distribution of their behaviorally associated edges. Specifically, for each bilateral seed nucleus, all edges that connect to the rest of the brain were correlated with the residual behavioral score of age and sex, and edges that meet $p < 0.05$ were separated by the correlation sign. The positive and negative edge strengths of the participants were then calculated by adding the DMCF values within each sign, and the net nodal strength (positive minus negative) was residualized for age and sex and correlated with the residualized behavioral score to produce an overall node-level association index ($R_{\text{Total}}$), thus confirming the aggregate behavioral relevance of the seed node. Finally, to visualize how each key node integrates or segregates across the 15 Du-15 cortical networks, circular connectograms were generated by projecting the retained positive or negative edges connecting the bilateral seed to the 400 Schaefer cortical parcels (omitting subcortico-subcortical edges for visual clarity), with edge color saturation scaled by $|r|$.

#### A.23 Factor-level decomposition of the key-node associations

We reasoned that, compared to broad behavioral phenotypes, the latent factors extracted through factor analysis represent more independent and unitary behavioral dimensions; consequently, if behavior is grounded in brain dynamics, these purer constructs should map onto cleaner and more dissociable neural patterns. To test this in an exploratory analysis ($N = 779$), we correlated all DMCF edges incident to three primary seed nuclei (the posterior caudate (pCAU), the medial AMYgdala (mAMY) and the anterior HIPpocampus (aHIP)) with age- and sex-residualized factor scores after the procedure in Section A.22. Specifically, we examine verbal and visual intelligence (Factors 2 and 8) for pCAU–$g$; social distress, well-being, and extraversion/friendship (Factors 1, 4, and 5) for mAMY–extraversion; and processing speed and visual intelligence (Factors 3 and 8, reflecting mental exploration and imaginative construction) for aHIP–openness. Circular connectograms of the retained subcortico-cortical edges ($p < 0.05$, ordered by the Du-15 networks) were plotted alongside the whole-trait profiles ($N = 785$ for $g$; $N = 917$ for extraversion and openness) on a shared color scale $|r|$ per sign. The edge signs for negatively loaded factors (Factors 2, 3, 8, and 5) were inverted so that red consistently denotes higher cognitive ability or higher extraversion/friendship, while factor 1 (social distress) was plotted in its native direction to provide a mirror comparison with extraversion.

#### A.24 Replication of the analytical pipeline in the ISYB dataset

To evaluate the generalizability of our findings in the independent ISYB cohort, we first derived a latent cognitive dimension because the ISYB protocol did not include the extensive cognitive battery used to estimate general intelligence ($g$) in the HCP dataset. Following the factor-analytic procedure described in Section A.19, six age- and sex-residualized behavioral indicators available in ISYB ($N = 165$; NEO agreeableness, extraversion, openness, and conscientiousness, the Multidimensional Scale of Perceived Social Support (MSPSS), and the Symbol Digit Modalities Test (SDMT)) were submitted to maximum-likelihood factor analysis with varimax rotation, yielding two latent factors with eigenvalues $> 1$ (explaining 31.7% of the total variance). Factor 2, which captured variance in SDMT cognitive performance ($r = -0.30$), served as the cognitive target for replication ($N = 146$ with neuroimaging data; lower scores reflect better cognitive performance), alongside the five NEO personality trait scores ($N = 162$). The Whole-brain DMCF profiles (93,096 edges) were then computed as detailed in Section A.18 by partitioning each participant's resting-state scan into $T = 9$ non-overlapping temporal layers of 20 time points. Using the same analytical framework established in the HCP cohort, we performed both a whole-brain association analysis (Section A.21; summarizing the top 500 positively and top 500 negatively correlated edges into age and sex-adjusted participant-level net co-occurrence frequency features) and a targeted subcortical key-node analysis (Section A.22). Specifically, we evaluated the primary positive and negative subcortical hubs identified in HCP—the pCAU and THA-DP for cognition (Factor 2), pCAU and THA-DA for agreeableness, mAMY and pPUT for extraversion, and aHIP for openness—quantifying the associations of their net nodal frequency features with behavior and visualizing their subcortico-cortical co-occurrence profiles ($p < 0.05$, with Factor 2 edge signs inverted so that red consistently indicates higher cognitive performance).

### Appendix B Supplementary Results

#### B.1 Interpretation of the Flexibility Map

The spatial distribution of the general flexibility revealed a pronounced boundary-specific organization of the functional architecture of the brain. Regions with high flexibility were located primarily at the topological boundaries where multiple functional systems converge, with particularly high values observed around the Sylvian fissure (Fig. B1a,d). This perisylvian zone interfaces with multiple functional systems—including the auditory (AUD), somatomotor (SMOT), language (LANG), action-mode (AMN), and default (DN) networks—and its elevated overall flexibility indicates frequent involvement in cross-system community reconfiguration, suggesting a potential role in functional-state transitions and inter-system information exchange. In contrast, the default network core subsystem (DN-B) exhibited relatively low flexibility. The core regions of DN-B, including the angular gyrus and posterior cingulate cortex (PCC), as well as the core regions of sensory and motor systems such as VISual (VIS) and somatomotor (SMOT) networks, showed relatively stable community affiliations in time windows, suggesting a more specialized functional organization that may support the stability of core functional processes.

Building on these findings, we further decomposed overall flexibility into disjoint and cohesive components across the cortex (Fig. B1b,c). Most cortical regions exhibited relatively low disjoint flexibility, with only a small subset showing a greater capacity for independent community adjustment. These regions were again concentrated around the Sylvian fissure, as well as the cingulate sulcus, and were predominantly located within or surrounding the AMN, with a smaller proportion located in the visual networks (VIS). Because the Sylvian fissure and cingulate sulcus lie at the confluence of the primary somatomotor networks (SMOT-A and SMOT-B), the AMN, and the premotor-posterior parietal rostral network (PM-PPr), this distribution reflects dense dynamic functional integration across these systems, forming a

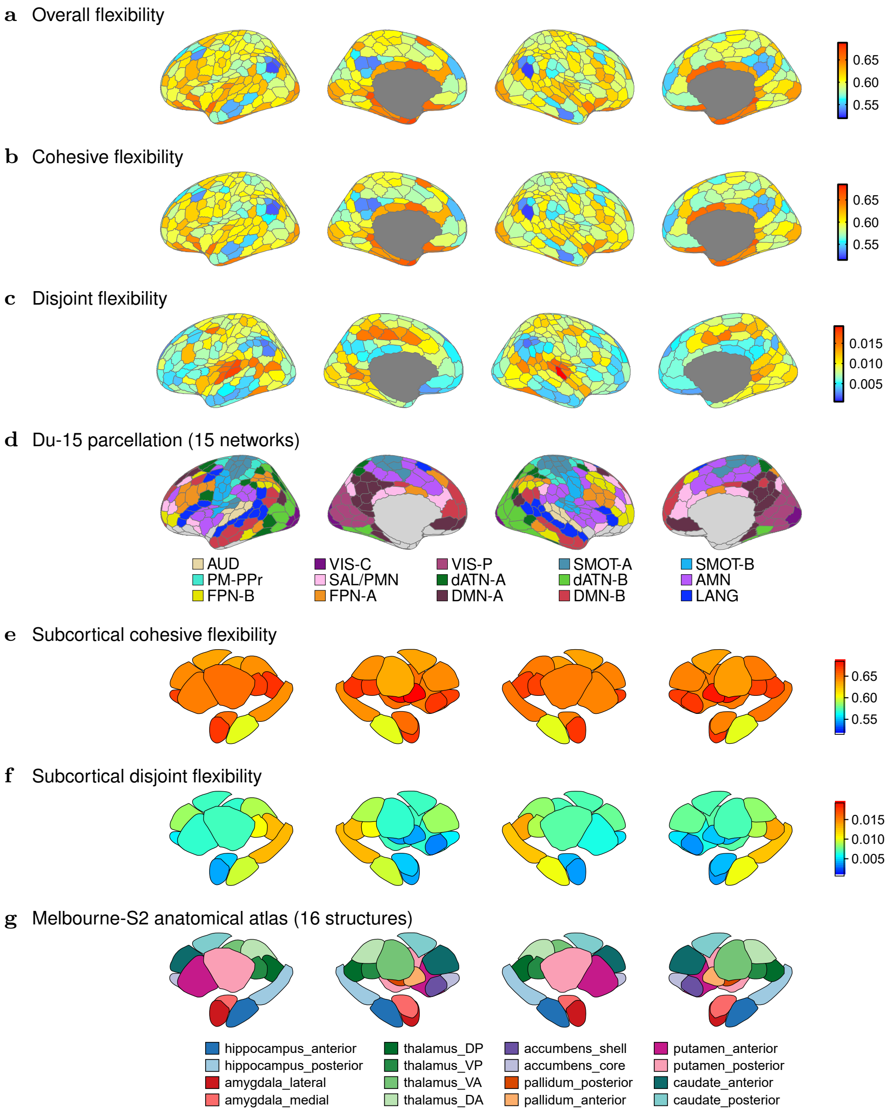


**Fig. B1 Flexibility of cortical and subcortical nodes relative to the Du-15 parcellation and the Tian atlas.** Each row shows, from left to right, the left lateral, left medial, right lateral and right medial views. **a**, Overall, **b**, cohesive and **c**, disjoint flexibility of the 400 cortical parcels; cohesive and disjoint flexibility sum to overall flexibility. **d**, The same parcels coloured by the Du-15 fifteen-network parcellation (grey, unassigned parcels and medial wall), with the network legend below the row. **e**, Subcortical cohesive and **f**, disjoint flexibility of the 32 Tian parcels, on the same colour scales as **b** and **c** (subcortical overall flexibility is shown in Fig. 2a). **g**, The Tian atlas underlying **e** and **f**, with its 16 nuclei-grouped structures labelled below the row and coloured by nucleus group. High overall and cohesive flexibility concentrates at network boundaries, whereas high disjoint flexibility is confined to a few parcels around the Sylvian fissure and the cingulate sulcus.

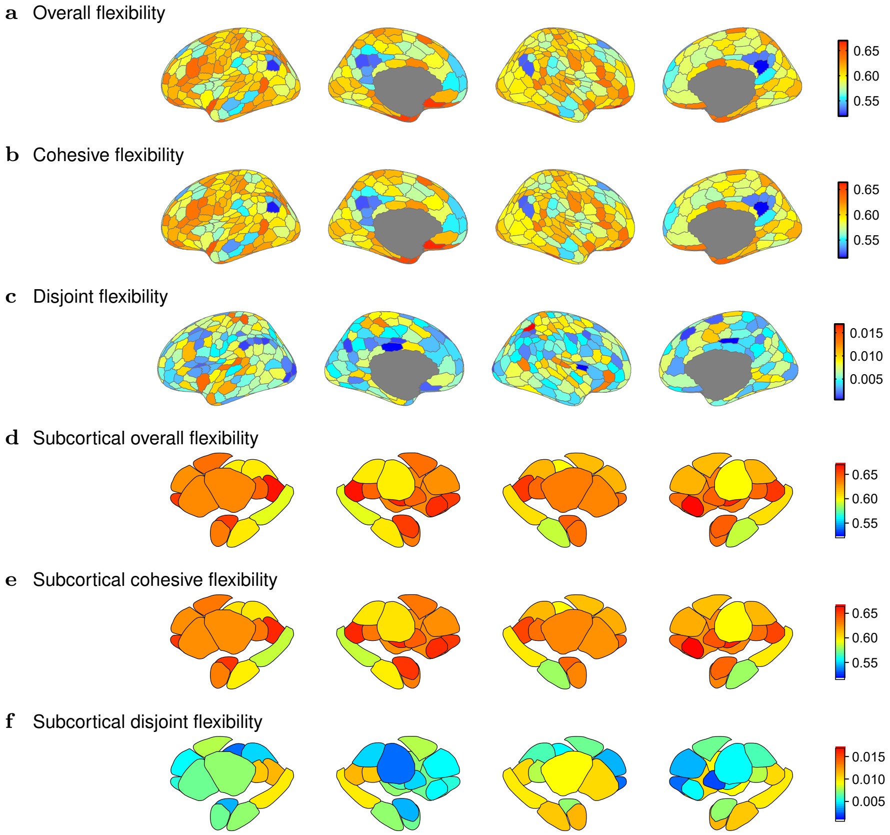


**Fig. B2 Cross-dataset replication of the flexibility topography in the independent ISYB cohort.** The layout follows Fig. B1, showing, from left to right, the left lateral, left medial, right lateral, and right medial views. **a**, Overall, **b**, cohesive, and **c**, disjoint flexibility of the 400 cortical parcels; **d**, overall, **e**, cohesive, and **f**, disjoint flexibility of the 32 Tian subcortical parcels, plotted on the same colour scales as **a**–**c**. Although colour scales span the specific value range of the ISYB cohort, the spatial topography across both cortical and subcortical structures closely replicates the HCP patterns (Fig. B1), consistently localizing high overall and cohesive flexibility to network boundaries and high disjoint flexibility to regions around the Sylvian fissure and cingulate sulcus.

bottom-up information-transmission framework extending from basic sensory systems toward action-related systems. The high disjoint flexibility of these regions suggests that a small number of nodes with strong local dynamic adjustment capacity may contribute to the initial stage of information transmission. In contrast, cohesive flexibility showed a more spatially distributed pattern, with higher values primarily observed in PM-PPr, frontoparietal network A (FPN-A), and parts of the central visual network

(VIS-C). Its overall spatial distribution was broadly consistent with that of overall flexibility, indicating that most nodes tend to participate in community reconfiguration in a cohesive manner and may subsequently contribute to broader information transmission and functional adjustment.

Extending this analysis to the subcortical parcels (Fig. B1e–g), subcortical nodes were on average more flexible than cortical nodes, in line with previous reports that subcortical structures are among the most flexible regions in the brain [115]. Specifically, the highest overall flexibility in the brain was observed in the pallidum and the dorsoposterior thalamus (THA-DP). As in the cortex, subcortical flexibility was driven almost entirely by cohesive rather than disjoint switching, with the highest cohesive values observed in the pallidum, THA-DP, lateral amygdala, and nucleus accumbens. This pattern indicates that subcortical nodes seldom change community affiliation in isolation, but instead reconfigure together within distributed cortico-striato-thalamo-cortical circuits [16, 53, 116]. Within this predominantly cohesive regime, however, THA-DP and the posterior hippocampus (pHIP) stood out for their relatively elevated disjoint flexibility (Fig. B1f). Because the posterior thalamus and posterior hippocampus interface extensively with cortical attentional, sensorimotor, and posteromedial systems [117–119], their elevated disjoint flexibility raises the possibility that these two structures represent putative boundary zones within the less-characterized functional parcellations of the subcortex, potentially serving as local initiation sites for subcortico-cortical reconfiguration [120].

Taken together, across both cortical and subcortical structures, disjoint flexibility captures the local adjustment capacity of a small number of network-boundary nodes, whereas cohesive flexibility reflects coordinated community reconfiguration involving a larger population of nodes. Their complementary spatial distributions suggest that brain network dynamics may follow a “leading” mechanism, whereby a small number of boundary nodes with high disjoint flexibility may undergo functional adjustment first and serve as early sites of information exchange, subsequently engaging broader populations of nodes in coordinated functional changes and thereby contributing to the dynamic cycling of the brain as a whole.

To evaluate whether this cortical and subcortical flexibility topography is consistent across datasets, we replicated the node-level analysis in the independent ISYB cohort ($n = 215$). Despite substantial differences in sample demographics and temporal sampling protocols (9 windows of 20 volumes in ISYB versus 20 windows of 60 volumes in HCP), the spatial patterns of all three flexibility measures were robustly reproduced across both cohorts (Fig. B2). Specifically, parcel-wise values in the ISYB dataset strongly correlated with the corresponding HCP maps for both cortical ($r = 0.77$, 0.77, and 0.41 for overall, cohesive, and disjoint flexibility, respectively) and subcortical ($r = 0.76$, 0.79, and 0.55) structures. Notably, the relatively moderate correlation for disjoint flexibility reflects its highly sparse spatial distribution—being driven by only a few localized boundary parcels—which naturally renders it more sensitive to shorter scan durations. Crucially, all core organizational features were consistently preserved across datasets: subcortical nodes remained on average more flexible than cortical nodes, overall and cohesive flexibility reliably concentrated along functional network boundaries, and elevated disjoint flexibility remained selectively anchored to the perisylvian cortex, cingulate sulcus, and posterior subcortex. This strong cross-dataset convergence demonstrates that the boundary-specific flexibility organization represents an intrinsic, generalizable feature of human brain dynamics rather than an artifact of a specific cohort or acquisition protocol.

#### B.2 Three-tier network model

To characterize the spatial organization of dynamic community co-occurrence across functional networks, we first examined community co-occurrence connections at the group level. We averaged the co-occurrence matrices across participants and constructed the group-level network by retaining edges in descending order of co-occurrence probability until every Du-15 functional network had at least one outward connection (1,057 edges; Fig. B3a), so as to capture the predominant community co-occurrence connections between functional networks. Each connection represents the frequency with which two brain regions were assigned to the same community during dynamic community detection.

We then grouped these high-co-occurrence connections according to the functional networks to which the corresponding brain regions belonged, allowing us to characterize shared community assignments between functional systems. Higher co-occurrence probabilities indicate that the corresponding brain regions were more frequently assigned to the same community across time windows, and therefore reflect recurrent and relatively stable organizational relationships during dynamic community reorganization. These high-co-occurrence connections revealed a relatively clear pathway of network reorganization. For example, the visual network (VIS) first showed strong community co-occurrence with the dorsal attention network-B (dATN-B), which in turn reorganized with the dorsal attention network-A (dATN-A) and extended towards the frontoparietal control networks-A and -B (FPN-A and FPN-B), ultimately reaching the default mode network (DMN), forming shared community assignments at a higher organizational level. This reorganization pathway reflects a progression from sensory information processing through attention- and action-related networks towards cognitive control and higher-order cross-modal cognition. Along this pathway, the relative positions of different networks within community reorganization exhibited a continuous hierarchical relationship, which could be further summarized into three interconnected functional tiers (sensory, attention/action, and regulatory), forming a hierarchical organization of network reorganization.

The third tier primarily comprised primary sensory and sensorimotor networks, including visual and somatomotor networks, and was characterized by strong within-network cohesion and relatively stable intranetwork integration. The second tier mainly included attention- and action-related networks, such as the dorsal attention, premotor and action-mode networks, which showed more pronounced cross-network shared community assignments and may support dynamic routing and integration of information across functional systems. The first tier primarily comprised transmodal regulatory networks, including the frontoparietal control and default mode networks, which exhibited widespread community co-occurrence with multiple functional systems, reflecting a higher degree of cross-system integration and potentially serving as key integrative nodes for higher-order cognition. Collectively, the high-co-occurrence connections formed a three-tier reorganization motif at the network level (Fig. B3c), extending progressively from primary sensory networks through attention- and action-related networks to transmodal regulatory networks. This pattern suggests that the fundamental integrative organization of dynamic brain networks follows a relatively stable hierarchical architecture.

Notably, some network co-occurrence relationships did not fully conform to this hierarchical organization. For example, the auditory network (AUD) bypassed the intermediate level and showed direct community co-occurrence with the language network (LANG). Previous work has shown that AUD and LANG are spatially adjacent in the lateral temporal cortex while remaining functionally distinguishable: AUD is primarily involved in sensory auditory processing, whereas LANG supports cross-modal language processing and responds to language presented in both auditory and visual modalities. Moreover, the spatial boundaries of LANG closely correspond to the transition between unimodal auditory functions and cross-modal language functions [121]. This organization indicates that auditory and cross-modal language processing are spatially adjacent in the lateral temporal cortex, forming a functional continuum from sensory processing towards higher-order language processing. Consistent with this organization, AUD exhibited relatively high dynamic flexibility and formed a community co-occurrence relationship with a functional system at a different hierarchical level, suggesting that its dynamic organization was not restricted to a single sensory system but instead involved substantial cross-community reorganization. Given the proximity of AUD to the spatial boundaries of multiple functional systems, such dynamic reorganization may provide a basis for its integration with distinct functional systems, resulting in a pronounced cross-level organizational pattern. Thus, the cross-level community co-occurrence between auditory and language networks not only reflects functional integration between sensory and higher-order cognitive systems but also provides macroscale evidence of direct dynamic coupling between auditory- and language-related functions.

The premotor network (PM-PPr) exhibited another distinct organizational pattern that deviated from the primary hierarchical pathway. It showed strong community co-occurrence primarily with sensorimotor networks, including SMOT-B and SMOT-A, but did not directly connect with higher-order control networks. Instead, its relationships with higher-order systems were mediated more prominently through the action-mode network, extending towards dATN-B and subsequently to higher-order control networks. Previous work has shown that the premotor cortex is involved in action selection and motor preparation and is closely associated with motor-related attentional processes [122]. This functional profile is consistent with the hierarchical position observed here, with PM-PPr situated closer to the preparation and early regulation of motor behavior and extending towards attentional and control systems through the action-mode network. Thus, PM-PPr occupies an intermediate position in the transition from motor-related networks towards attention and cognitive control systems.

In contrast to the networks described above, the salience network (SAL) exhibited strong cross-network community co-occurrence with the default mode network (DMN) and higher-order cognitive networks, including FPN-A and FPN-B, while also showing cross-hierarchical community co-occurrence with the action mode network (AMN). This pattern suggests an organizational structure that differs from the predominantly hierarchical, top-down transmission pathways observed across the main functional hierarchy. Previous studies have shown that the SAL is involved in detecting behaviorally salient information and can facilitate dynamic switching between the DMN and the central executive network, thereby regulating the allocation of attentional and cognitive control resources [3, 36]. In light of the observed community co-occurrence patterns, the cross-network dynamics involving the SAL span attentional, control, and higher-order cognitive systems. Its co-occurrence with the AMN and multiple higher-order cognitive networks may therefore reflect its involvement in information transmission across control and attentional hierarchies.

To test whether this three-tier organization generalized beyond the HCP cohort, we repeated the group-level network analysis in the ISYB cohort using the same edge-retention rule, which yielded 1,023 edges (Fig. B3b). Overall, the fundamental three-tier hierarchical framework was preserved. For example, along the sensorimotor pathway, the somatomotor networks (SMOT-A and SMOT-B) still connected first to the premotor (PM-PPr) and action-mode (AMN) networks in the intermediate tier before extending to the frontoparietal control networks (FPN-A and FPN-B) in the regulatory tier. However, two notable differences emerged in the sensory pathways: the outward connection of the auditory network (AUD) was directed to the AMN rather than to the language network (LANG), whereas the peripheral visual network (VIS-P) exhibited direct connections to the default network (DN-A). Given the differences between the ISYB and HCP cohorts in sample size, temporal sampling (9 versus 20 time windows per run), and acquisition protocols, these pathway-level variations may reflect either sampling variability or cohort-specific characteristics, without altering the overarching three-tier architecture.

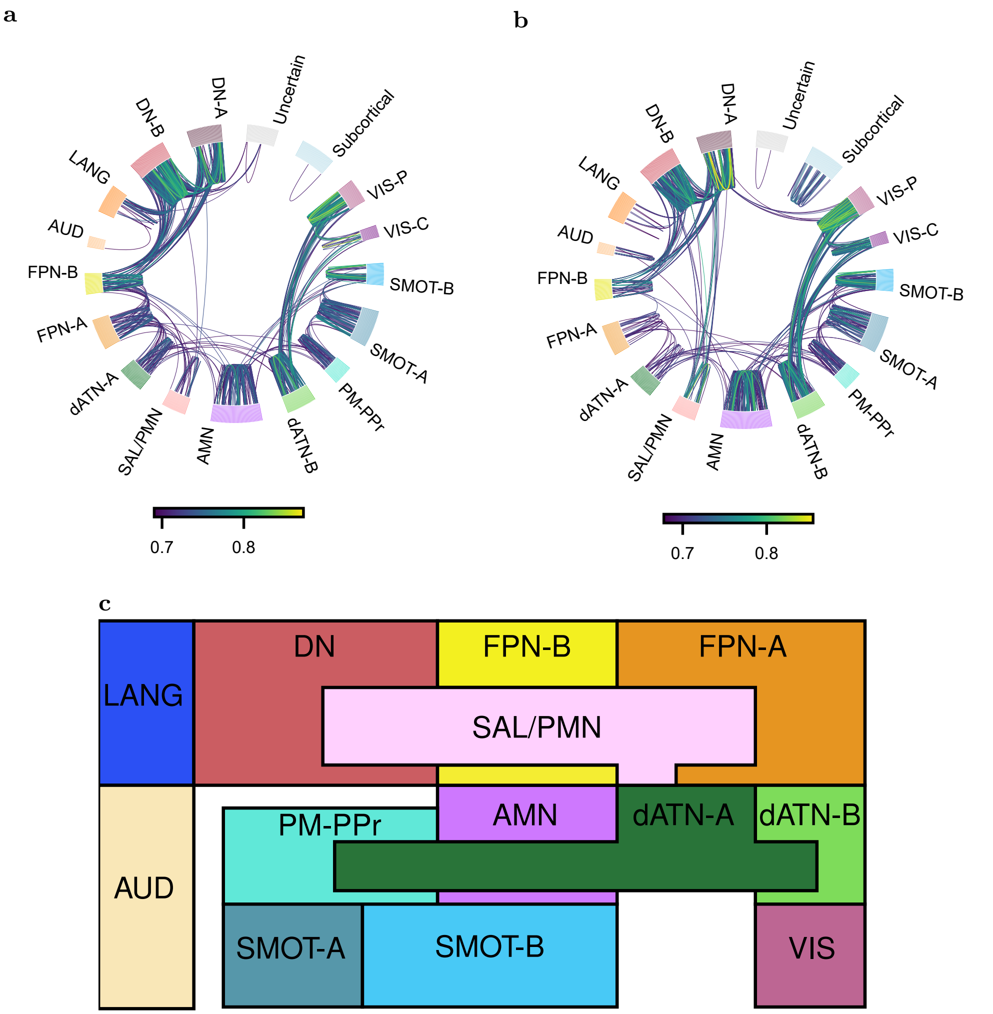


**Fig. B3 Group-level community co-occurrence networks in HCP and ISYB, and the three-tier organization. a**, HCP network. Edges were added in order of decreasing co-occurrence probability until every Du-15 functional network had at least one outward connection (1,057 edges). **b**, ISYB network. Applying the same edge-retention rule yielded 1,023 edges. While the overall three-tier hierarchy is preserved, AUD connects outward to AMN and VIS-P connects directly to DN-A. In **a** and **b**, nodes are arranged by Du-15 functional network around the ring, and edge colour denotes co-occurrence probability. **c**, Three-tier organization of dynamic community co-occurrence. Group-level co-occurrence connections summarize the progression from primary sensory networks through attention/action networks to transmodal regulatory networks.

#### B.3 Maximum-Overlap Community Map

In the two-community partition, the consensus map divided the cortex into two large systems with opposing spatial emphases (Fig. B4a). One community occupied the default-mode, control, and limbic territories, oriented toward internal processing; the other encompassed the somatomotor, dorsal-attention, visual, and salience regions, oriented toward the external environment. This inward/outward duplex mirrors the well-known antagonism between the default mode network (DMN) and the action mode network (AMN), which support internally and externally directed cognition, respectively [9, 24, 25]. We infer that this antagonism is not merely a static opposition, but a fundamental driver of continuous dynamic reconfiguration: as the balance between the two systems shifts over time, brain regions are repeatedly reassigned between the inward and outward poles, preserving this primary axis in the time-integrated static map. This robust duplex was consistently observed across both the HCP and ISYB cohorts, with the inward community capturing most default and control regions, and the outward community encompassing all somatomotor and dorsal-attention regions (Fig. B4a).

In the three-community partition, this overarching two-system division persisted, but an intermediate community emerged between the inward and outward poles (Fig. B4b). Across both cohorts, this transitional community was predominantly composed of salience and control regions, alongside a subset of dorsal-attention areas. Its intermediate topological position aligns closely with the established role of the salience network in mediating switches between the DMN and central-executive systems [3, 36]; we therefore refer to it as a bridging network.

In the seven-community partition, the map resolved into functional modules whose spatial layout closely resembled the canonical Yeo-7 parcellation [2] (Fig. B4c). Evaluating the spatial alignment across non-limbic cortex, the consensus communities achieved high overlap rates of 90.16% in HCP and 89.10% in ISYB. This strong cross-cohort agreement indicates that the seven-community consensus map captures the canonical functional architecture of the brain, rather than reflecting a cohort-specific configuration.

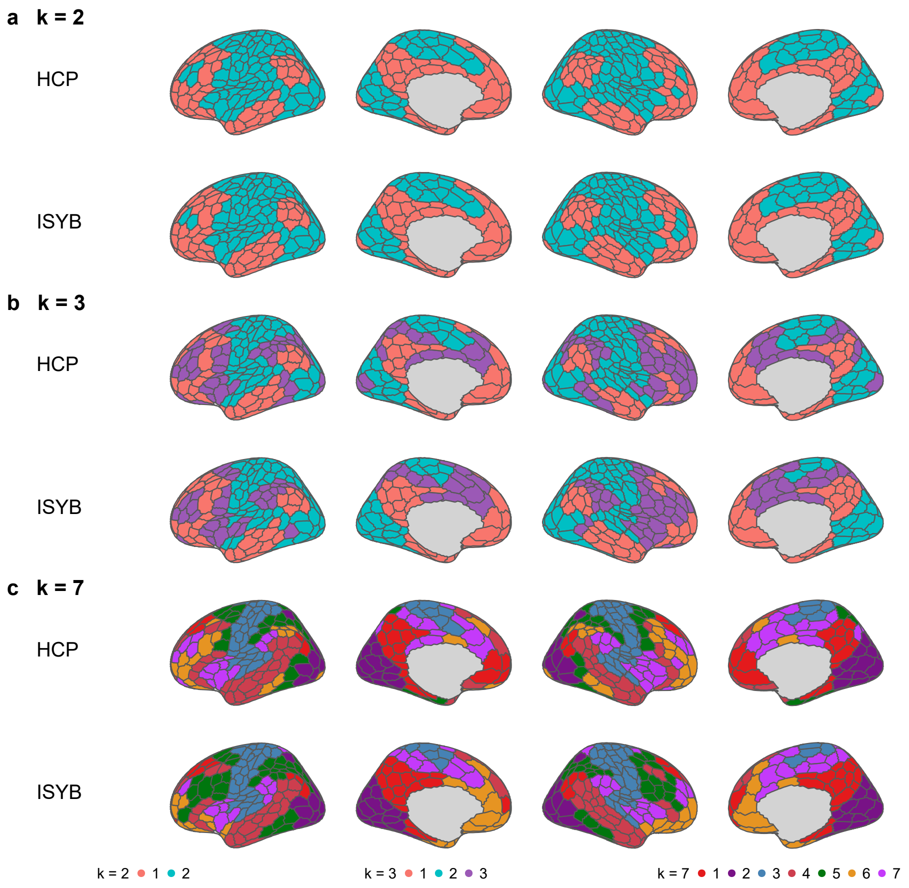


**Fig. B4 Maximum-overlap community organization in HCP and ISYB.** Each panel shows the group-level consensus partition on the 400-parcel Schaefer cortical surface in four views (left lateral, left medial, right lateral, right medial), with HCP above and ISYB below; colours denote consensus communities, and the legend for each $k$ is shown at the bottom. **a**, $k = 2$. **b**, $k = 3$. **c**, $k = 7$. At $k = 2$ the partition separates an inward-facing default/control system from an outward-facing sensorimotor/attention system; at $k = 3$ a salience-control bridging community emerges between them; at $k = 7$ the partition approaches the Yeo-7 organization, with best-two overlap rates of 90.16% (HCP) and 89.10% (ISYB) over the 376 non-limbic cortical parcels.

#### B.4 Detailed Results of the Three Subcortical Routing Modes

The basal mode is anchored by the ventral posterior thalamus (THA-VP), whose co-occurrence pattern map is illustrated in Fig. B5. High co-occurrence probabilities with THA-VP were specifically enriched within primary and early sensory cortices, well above the whole-brain average. Specifically, these regions encompassed the bilateral postcentral gyrus (primary somatosensory cortex), pericalcarine cortex and lingual gyrus (early visual cortex), and bilateral superior temporal and Heschl's gyri (primary and early auditory cortices). Conversely, core nodes of the default mode network (e.g., medial prefrontal and posterior cingulate cortices) and higher-order association cortices (e.g., anterior lateral temporal cortex) exhibited minimal co-occurrence (with ventromedial regions masked due to signal quality constraints). This distinct topological profile indicates that THA-VP selectively forms highly synchronized co-activation patterns with specific visual, auditory, and somatosensory networks.

Two non-mutually exclusive mechanisms may underlie this phenomenon: (i) during the dynamic self-organization of brain networks, THA-VP may function beyond a unimodal somatosensory relay, participating deeply in bottom-up, cross-modal multisensory integration and temporal synchronization [13, 14, 123]; and (ii) given the close anatomical proximity of the ventral posterior thalamus to the medial and lateral geniculate bodies, this co-activation may reflect the concerted operation of the entire posterior thalamic sensory complex in processing multisensory inputs [106, 124].

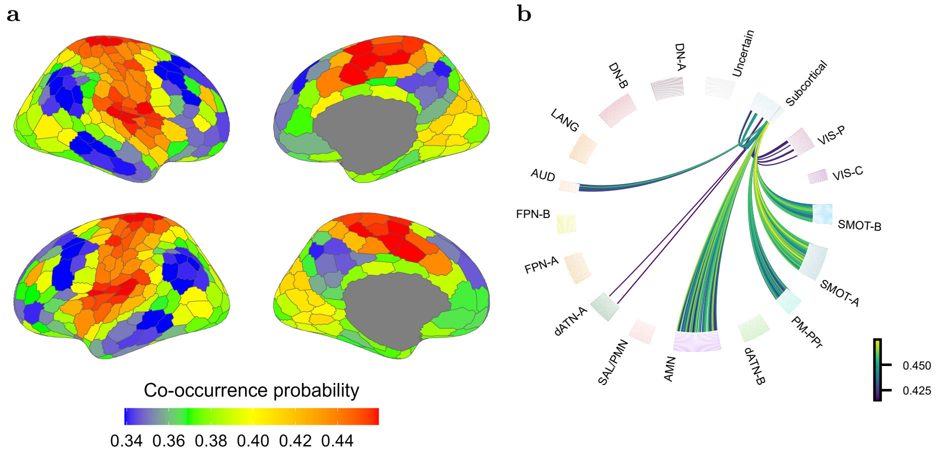


**Fig. B5 Basal mode.** **a**, Co-occurrence map of the ventral posterior thalamus (THA-VP, red), averaged over the left and right THA-VP. **b**, Circular connectivity plot of the bilateral THA-VP (strongest 30% of connections; chord colour: co-occurrence probability; the fused bilateral THA-VP node in the colour of the basal routing mode). The ring follows the network order of the main text.

The regulatory mode is anchored by a distributed cortico-striato-thalamic circuit, encompassing associative thalamic subdivisions (THA-VA, THA-DA, THA-DP), the caudate (aCAU, pCAU), the putamen (aPUT, pPUT), the nucleus accumbens (NAc-shell, NAc-core), and the pallidum (aGP, pGP). At the macroscale, this mode predominantly integrates tier 1 and tier 2 cortical systems.

Within this mode, individual subcortical nodes exhibit specialized cross-network coupling profiles that mirror their complementary computational roles in the canonical cortico-basal ganglia-thalamic architecture (Fig. B6). First, the nucleus accumbens (NAc) couples the salience network (SAL), default mode network, and FPN-B. Consistent with its role as a central reward hub that assigns motivational salience [42], this tripartite coupling suggests that the NAc acts as an affective-motivational interface, translating internal self-referential states and value signals into salience-driven engagement of executive control. Second, the caudate selectively bridges the SAL, FPN, and DN-B. This profile aligns with its established involvement in goal-directed behavior and the flexible updating of cognitive strategies [43], indicating that the caudate coordinates executive control circuits and specific default mode subsystems to dynamically adapt internal strategies to salient environmental demands. Third, the putamen and posterior pallidum (pGP) selectively integrate across the action mode network and SAL. This pattern reflects the role of the putamen in sensorimotor integration and habitual control [44] alongside the function of the pallidum as a basal ganglia output gate for action selection [45]: by directly linking salience detection to motor execution systems, these nodes facilitate rapid, automatized behavioral outputs triggered by salient cues. Finally, the associative thalamic subdivisions and anterior pallidum (aGP) display the most extensive integration profiles, spanning all of the aforementioned cortical systems, with the dorsoposterior thalamus (THA-DP) additionally integrating the visual network. This broad convergence underscores their role as integrative hubs [14] that synthesize parallel motivational, cognitive, and motor streams from the basal ganglia while incorporating bottom-up visuospatial inputs via the posterior thalamus [117]. Together, the convergence of these nuclei within a single routing mode points to a continuous processing stream that progresses from internal valuation and motivational salience, through flexible cognitive adaptation, to the gating of external behavioral output.

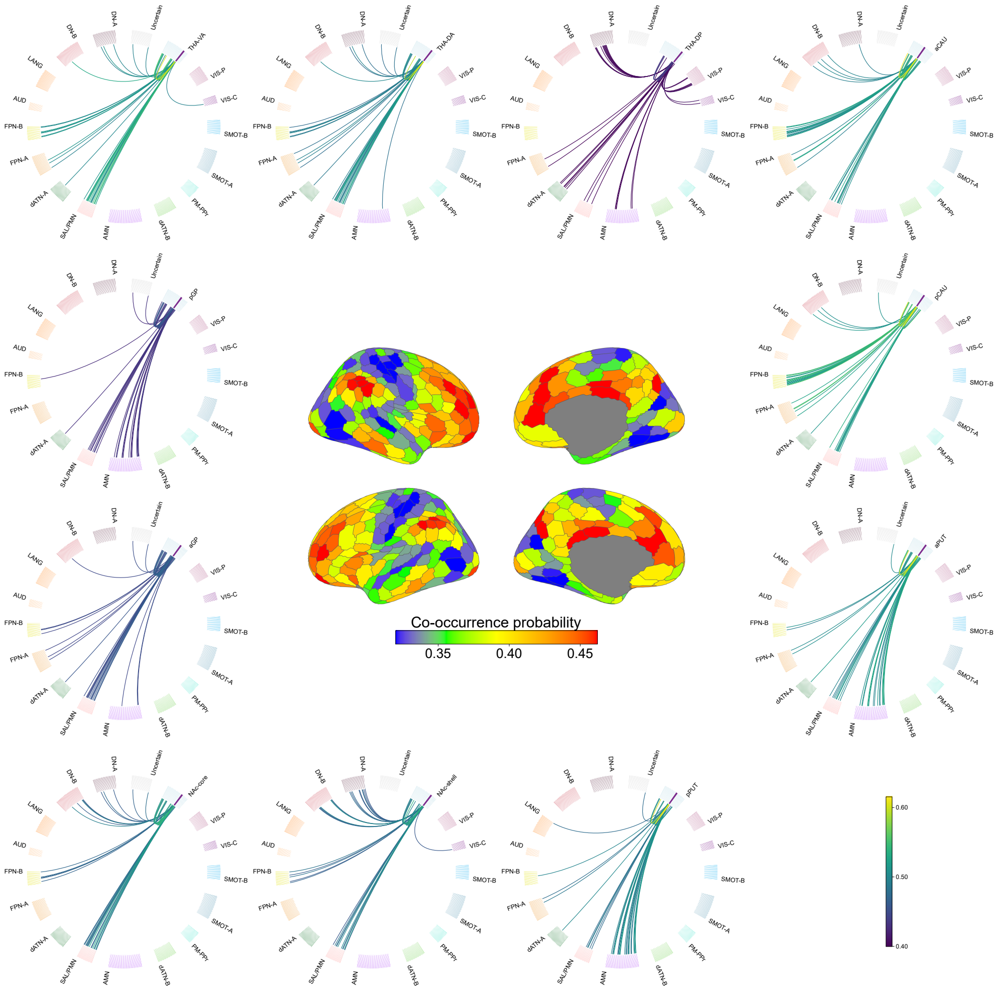


**Fig. B6 The regulatory mode across its eleven constituent nuclei.** The eleven nuclei of the regulatory mode (THA-VA, THA-DA, THA-DP, aCAU, pCAU, aPUT, pPUT, NAc-shell, NAc-core, aGP, pGP). The central panel is their group-average co-occurrence map on four cortical views (left lateral, left medial, right lateral, right medial). The eleven surrounding rings show each nucleus separately, clockwise from the upper left in the order above, with all 432 parcels in the network order of the main text; the focal nucleus is drawn in the regulatory-mode colour (purple) and the chords give its strongest 10% of connections (43 per nucleus, left and right instances fused). The bar below the central map gives its co-occurrence probability (0.32–0.46, the 5th–95th percentile of the 400 cortical parcels; blue to red); the vertical bar at the lower right is shared by all eleven rings and gives their chord co-occurrence probability (0.40–0.62; viridis).

The fast mode comprises the hippocampus (aHIP, pHIP) and the amygdala (mAMY, lAMY) (Fig. B7). At the macroscale, this mode predominantly integrates across the default mode network and the motor network, whereas the posterior hippocampus (pHIP) specifically couples the default mode network with FPN-B. This topological feature not only reflects the functional division between the anterior and posterior hippocampus [118], but also provides a critical transitional outlet for the fast mode to dynamically switch into the other two routing modes. Canonically, the hippocampus governs memory, spatial navigation, and episodic reconstruction and imagination [60], whereas the amygdala mediates emotional processing and the formation of emotional memories [46]. Although these established functions largely center on self-referential and mnemonic processing, our community co-occurrence patterns reveal an additional role for this mode: an emotion-driven rapid pathway from internal states to external action. Together, this mode is poised to drive rapid network reorganization in response to environmental crises, providing an intrinsic impetus for the dynamic reconfiguration of functional networks.

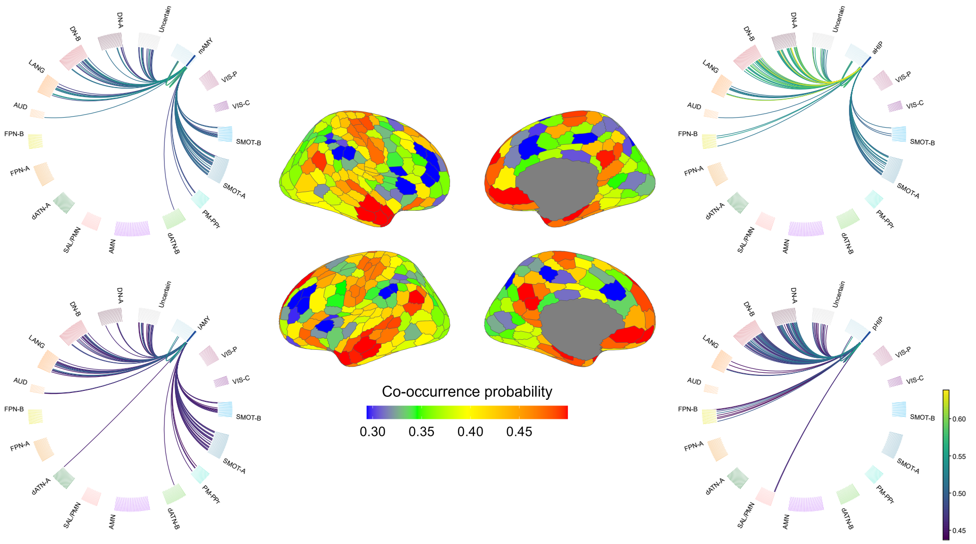


**Fig. B7 Fast mode.** The four nuclei of the fast mode (blue): the amygdala, medial (mAMY, upper left) and lateral (lAMY, lower left), and the hippocampus, anterior (aHIP, upper right) and posterior (pHIP, lower right). The central panel is the mean co-occurrence map of the four nuclei on four cortical views (left lateral, left medial, right lateral, right medial), dominated by coupling with the default mode and motor networks; the horizontal bar below it gives its co-occurrence probability (0.29–0.50, the 5th–95th percentile of the 400 cortical parcels; blue to red). The four rings show each nucleus separately, with all 432 parcels in the network order of the main text; the posterior hippocampus is distinguished by its coupling between the default mode network and FPN-B. The focal nucleus is drawn in the fast-mode colour (blue) and the chords give its strongest 20% of connections (86 per nucleus, left and right instances fused). The vertical bar is shared by all four rings and gives their chord co-occurrence probability (0.44–0.64; viridis).

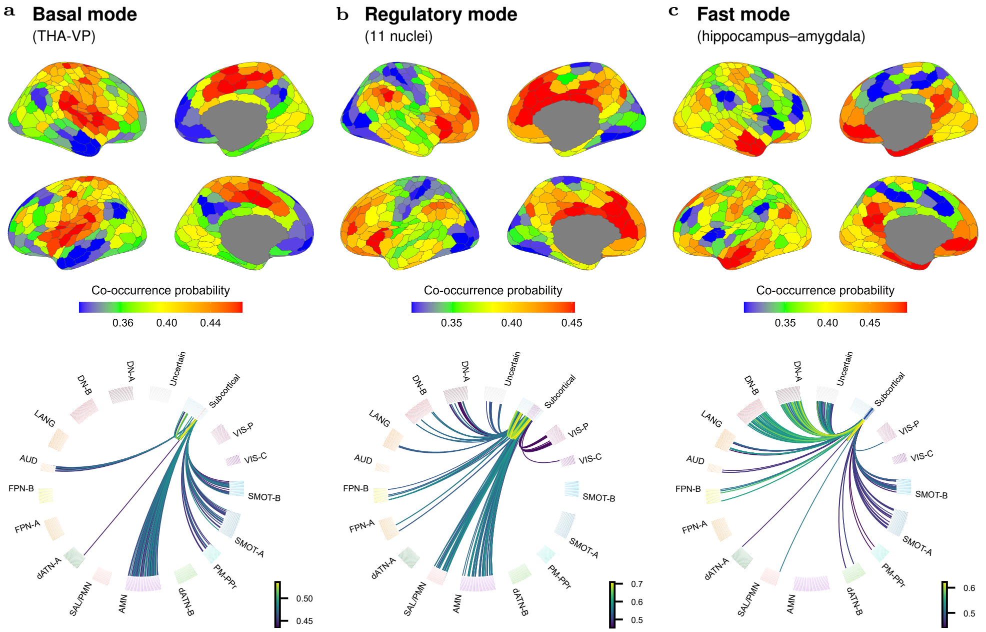


**Fig. B8 Cross-dataset replication of the three subcortical routing modes in the ISYB cohort.** **a**–**c**, Basal (**a**), regulatory (**b**), and fast (**c**) modes in the ISYB cohort ($n = 215$), derived using the same anatomical definitions and pipeline as Figs. B5–B7. Upper row, four-view cortical maps of the mean co-occurrence probability between the nuclei of each mode and the 400 Schaefer parcels; each colour bar spans the 5th–95th percentile of its corresponding cortical values (0.32–0.47, 0.32–0.45, and 0.30–0.49; blue to red). Lower row, circular connectivity plots (chord colour: co-occurrence probability). **a**, Bilateral THA-VP fused into a single node (431 nodes), showing the strongest 20% of connections per side (97 chords). **b**, The eleven striato-thalamo-pallidal nuclei (432 nodes), showing the strongest 5% per nucleus (226 chords). **c**, The four hippocampal and amygdalar nuclei (428 nodes after fusing left–right counterparts), showing the strongest 15% per nucleus (245 chords). Rings follow the network order of the main text. Stricter percentile thresholds than in Figs. B5–B7 are applied purely for visualization to maintain comparable chord densities in the smaller ISYB rings. Although absolute co-occurrence probabilities lie on a coarser grid due to fewer temporal windows in ISYB (9 versus 20 in HCP), the spatial coupling patterns and network-level routing topologies are consistently preserved across cohorts.

To evaluate whether these three subcortical routing modes are consistent across datasets and populations, we replicated the mode-level analysis in the independent Chinese ISYB cohort ($n = 215$; 9 windows of 20 volumes; Appendix A.24), applying the identical anatomical definitions, 432-node Tian–Schaefer parcellation, and percentile-based edge-retention framework (Fig. B8). Despite differences in cohort demographics and temporal sampling protocols, the cortical coupling maps were reproduced with remarkable fidelity: parcel-wise Pearson correlations between the HCP and ISYB maps across the 400 cortical parcels reached $r = 0.880$ for the basal mode, $r = 0.891$ for the regulatory mode, and $r = 0.866$ for the fast mode. Furthermore, circular connectivity profiles in the ISYB cohort consistently recovered the same canonical network-level coupling motifs—with THA-VP selectively anchored to primary sensory cortices, the striato-thalamo-pallidal circuit distributed across salience, control, and action-related systems, and the hippocampal–amygdalar complex bridging the default mode and motor networks. Thus, although fewer temporal windows in ISYB place absolute co-occurrence probabilities on a coarser numerical grid, the relative coupling hierarchy and macroscale routing topology remain highly invariant. This strong cross-dataset convergence demonstrates that the three subcortical routing modes reflect fundamental, generalizable principles of human cortico-subcortical organization.

#### B.5 Behavioral target selection and factor structure

Initial connectome-based predictive screening coupled with manual curation (Appendix A) identified 29 candidate behavioral and physiological measures that showed significant out-of-fold predictability from the DMCF ($r = 0.075$–$0.258$, uncorrected $p < 0.05$; see Table B1 for full statistics). These measures represent a diverse profile spanning cognition, emotion, motor function, personality, task performance, and physiology.

**Table B1 Pooled out-of-fold predictability of 29 measures.**

| Behavior | $N$ | $r$ | $p$ |
|---|---|---|---|
| *Cognition (9)* | | | |
| PicVocab_Unadj | 711 | 0.258 | $2.9 \times 10^{-12}$ |
| ReadEng_Unadj | 711 | 0.206 | $2.9 \times 10^{-8}$ |
| PMAT24_A_CR | 711 | 0.190 | $3.5 \times 10^{-7}$ |
| PicSeq_Unadj | 711 | 0.169 | $6.2 \times 10^{-6}$ |
| DDisc_AUC_200 | 711 | 0.152 | $4.8 \times 10^{-5}$ |
| ProcSpeed_Unadj | 711 | 0.141 | $1.6 \times 10^{-4}$ |
| CardSort_Unadj | 711 | 0.133 | $3.9 \times 10^{-4}$ |
| IWRD_TOT | 711 | 0.115 | 0.002 |
| VSPLOT_TC | 711 | 0.084 | 0.025 |
| *Emotion (8)* | | | |
| AngAggr_Unadj | 710 | 0.180 | $1.4 \times 10^{-6}$ |
| Loneliness_Unadj | 710 | 0.118 | 0.002 |
| MeanPurp_Unadj | 710 | 0.102 | 0.007 |
| LifeSatisf_Unadj | 710 | 0.096 | 0.011 |
| PercReject_Unadj | 710 | 0.095 | 0.011 |
| Friendship_Unadj | 710 | 0.089 | 0.018 |
| FearAffect_Unadj | 710 | 0.083 | 0.026 |
| SelfEff_Unadj | 710 | 0.082 | 0.029 |
| *Motor (3)* | | | |
| Endurance_Unadj | 711 | 0.161 | $1.6 \times 10^{-5}$ |
| Strength_Unadj | 710 | 0.117 | 0.002 |
| GaitSpeed_Comp | 711 | 0.108 | 0.004 |
| *Personality (3)* | | | |
| NEOFAC_A | 711 | 0.108 | 0.004 |
| NEOFAC_E | 711 | 0.090 | 0.016 |
| NEOFAC_O | 711 | 0.075 | 0.046 |
| *In-Scanner Task Performance (4)* | | | |
| Relational_Task_Match_Median_RT[1] | 703 | 0.150 | $6.3 \times 10^{-5}$ |
| WM_Task_2bk_Tool_Median_RT_Target[1] | 690 | 0.133 | $4.8 \times 10^{-4}$ |
| Language_Task_Story_Median_RT[1] | 709 | 0.119 | 0.002 |
| Emotion_Task_Shape_Acc | 707 | 0.078 | 0.037 |
| *Physiology (2)* | | | |
| BPDiastolic[1] | 709 | 0.218 | $4.5 \times 10^{-9}$ |
| PSQI_AmtSleep[1] | 711 | 0.185 | $6.8 \times 10^{-7}$ |

$N$, participants with complete data; $r$, Pearson correlation between pooled out-of-fold predictions and the age- and sex-residualized measure (10-fold cross-validated CPM; 500 positive and 500 negative edges per fold); $p$, two-sided test on $r$, uncorrected.

[1]Measures added in the second screening run.

**Table B2 Pooled out-of-fold prediction performance of the 16 target phenotypes.**

| Target | Type | $N$ | $R_{\text{OOF}}$ | $p_{\text{perm}}$ | $q_{\text{BH}}$ |
|---|---|---|---|---|---|
| *Cognition (7)* | | | | | |
| Verbal Intelligence | Factor | 828 | 0.457 | 0.001 | 0.003 |
| Delayed Gratification | Independent | 1027 | 0.222 | 0.001 | 0.002 |
| Picture Sequence Memory | Independent | 1030 | 0.213 | 0.001 | 0.002 |
| Processing Speed | Factor | 828 | 0.190 | 0.001 | 0.003 |
| Story Task Reaction Time | Independent | 996 | 0.118 | 0.004 | 0.005 |
| Visual Intelligence | Factor | 828 | 0.093 | 0.020 | 0.023 |
| Verbal Episodic Memory | Independent | 1027 | 0.088 | 0.019 | 0.022 |
| *Personality/social (5)* | | | | | |
| Extraversion and Friendship | Factor | 828 | 0.119 | 0.008 | 0.011 |
| Social Distress | Factor | 828 | 0.117 | 0.003 | 0.006 |
| Well-being | Factor | 828 | 0.107 | 0.007 | 0.011 |
| Aggression | Factor | 828 | 0.074 | 0.048 | 0.048 |
| Expression Recognition | Independent | 991 | 0.063 | 0.065 | 0.065 |
| *Physiology/motor (4)* | | | | | |
| Physical Fitness | Factor | 828 | 0.193 | 0.001 | 0.003 |
| Sleep | Independent | 1031 | 0.190 | 0.001 | 0.002 |
| Blood Pressure | Independent | 1018 | 0.182 | 0.001 | 0.002 |
| Strength | Independent | 1030 | 0.137 | 0.001 | 0.002 |

$N$, participants with complete data; $R_{\text{OOF}}$, Pearson correlation between pooled out-of-fold predictions (nested 10-fold outer cross-validation with 5-fold inner tuning of the ridge penalty) and the age- and sex-residualized target; $p_{\text{perm}}$, one-sided permutation test (1,000 permutations); $q_{\text{BH}}$, Benjamini–Hochberg false discovery rate computed separately within the eight latent factors and within the eight independent components. Factors are the eight latent dimensions from the exploratory factor analysis; independent components are the eight measures with $|\text{loading}| < 0.30$ on every factor.

To uncover the latent structure among these measures, we submitted them to an exploratory factor analysis ($N = 868$), which extracted eight nearly orthogonal factors (maximal inter-factor $|r| = 0.18$) accounting for 34.9% of the total variance (Fig. B9). Twenty-one measures loaded strongly ($|\text{loading}| \geq 0.30$) onto at least one factor, whereas the remaining eight fell below this threshold across all factors and were retained as independent components. Together, these eight factors and eight independent components defined 16 behavioral targets organized into three broad domains (Fig. 3a; Table B2):

**Cognitive domain (seven targets).** Three factors captured core cognitive dimensions: *verbal intelligence* (Factor 2), anchored by vocabulary, reading, matrix

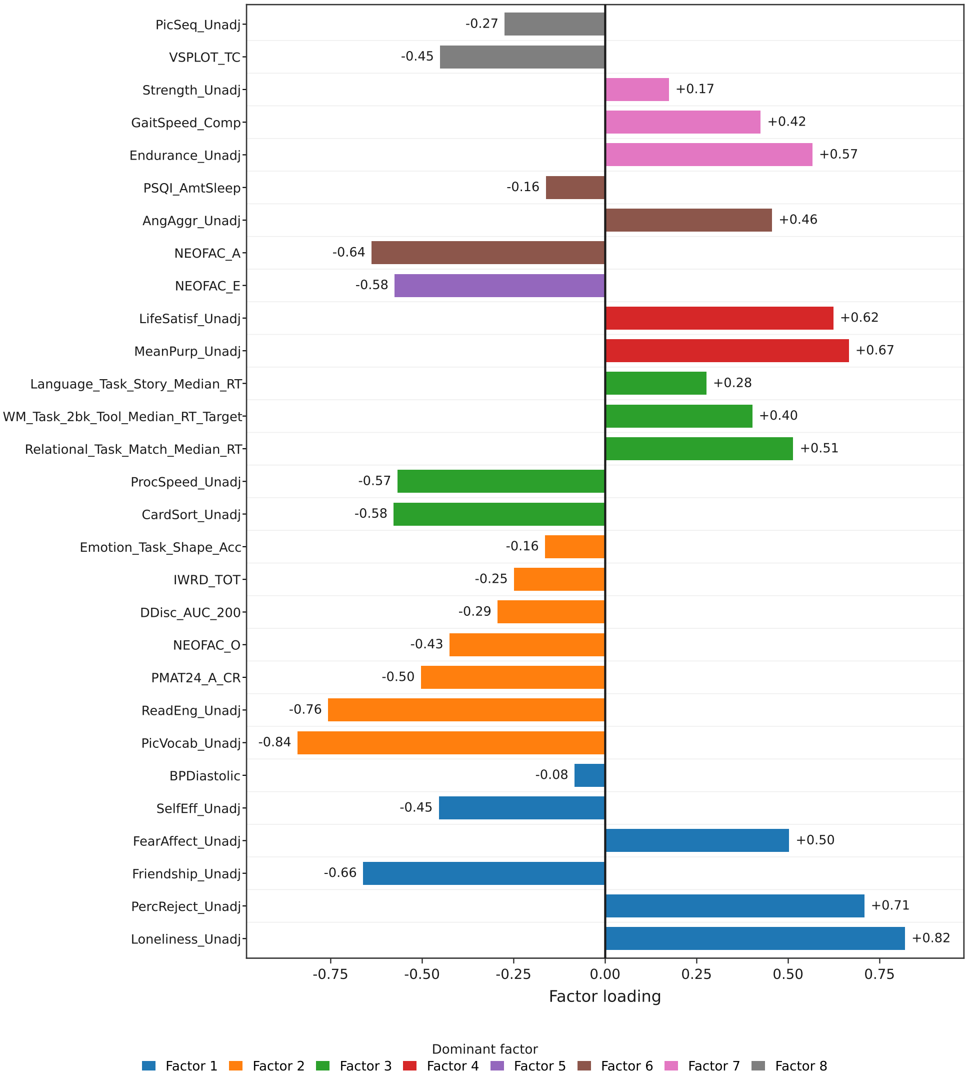


**Fig. B9 Dominant factor loading of each of the 29 measures.** For each measure the bar gives its signed loading on the factor on which its absolute loading is largest, and its colour gives that factor (Factor 1, social distress; Factor 2, verbal intelligence; Factor 3, processing speed; Factor 4, well-being; Factor 5, extraversion and friendship; Factor 6, aggression; Factor 7, physical fitness; Factor 8, visual intelligence). Factor 1 is at the bottom and Factor 8 at the top, and the members of each factor are ordered by absolute loading. Values at the bar ends are the loadings. The eight measures whose maximal absolute loading was below 0.30 are coloured by their largest loading but do not define any of the eight factors.

reasoning, and openness; *processing speed* (Factor 3), integrating card sorting and pattern comparison speed with reaction times from the relational-matching and working-memory tasks; and *visual intelligence* (Factor 8), driven primarily by visuospatial orientation performance. Four cognitive measures with sub-threshold loadings entered as independent components: picture sequence memory, delay discounting (AUC), story-task reaction time, and verbal episodic memory.

**Personality and socio-emotional domain (five targets).** Four factors summarized socio-emotional functioning: *social distress* (Factor 1), coupling loneliness, perceived rejection, and fear affect with reduced friendship and self-efficacy; *well-being* (Factor 4), defined by meaning in life and life satisfaction; *extraversion and friendship* (Factor 5), carried primarily by extraversion alongside secondary contributions from friendship and openness; and *aggression* (Factor 6), a bipolar axis opposing aggression against agreeableness. Emotion expression recognition entered as an independent component.

**Physiological and motor domain (four targets).** A single factor, *physical fitness* (Factor 7), linked physical endurance with gait speed, while grip strength, diastolic blood pressure, and sleep duration were retained as independent components.

#### B.6 Whole-brain connectograms of the associations with intelligence and personality

A key advantage of our edge-level framework is the ability to trace overall behavioral associations back to the specific inter-network integration features that drive them. Having established that the DMCF profile tracks a broad range of behaviors (Section B.5), we examined the large-scale network architecture underlying two higher-order constructs: general intelligence ($g$), derived from ten HCP cognitive measures, and personality, represented by the three NEO Five-Factor dimensions that passed screening (agreeableness, extraversion, and openness; Appendix A). Projecting the 500 most positively and 500 most negatively correlated edges onto the 432-node connectogram revealed robust net integration signatures across all four targets (Fig. B10a–d, left panels), with $g$ ($r = 0.424$, $p = 1.3 \times 10^{-35}$) and openness ($r = 0.372$, $p = 2.0 \times 10^{-31}$) exhibiting stronger associations than agreeableness ($r = 0.265$, $p = 3.4 \times 10^{-16}$) and extraversion ($r = 0.258$, $p = 2.2 \times 10^{-15}$).

We interpreted these connectograms through the lens of the three-tier cortical hierarchy[31]—spanning first-tier transmodal association networks (default and frontoparietal), second-tier attentional and action-mode networks, and third-tier primary sensorimotor and visual cortices. General intelligence ($g$) recapitulated the classic Parieto-Frontal Integration Theory (P-FIT)[125] while revealing a complementary pattern of hierarchical segregation (Fig. B10a, middle and right panels). Positive edges densely interconnected first- and second-tier executive and regulatory systems (FPN-A, SAL/PMN, AMN, and DN-A) and coupled FPN-A with somatomotor cortex, whereas negative edges strongly segregated the first-tier default network and AMN from second- and third-tier sensorimotor, language, and visual cortices. Together, these motifs indicate that higher intelligence is supported not only by strong integration across upper-tier control networks—facilitating deep cognitive strategy processing—but also by the functional insulation of these transmodal systems from lower-tier sensory interference.

Extending this hierarchical perspective to the three personality dimensions revealed distinct topological motifs for each trait (Fig. B10b–d, middle and right panels). Agreeableness was characterized by tight positive coupling within and between tiers—both between the first-tier default subsystems (DN-A–DN-B) and across second- and third-tier attentional and motor networks (dATN-B, PM-PPr, and SMOT-A/B)—paired with dense cross-tier negative edges that segregated the default network from the action-mode (AMN) and attention systems. This architecture suggests that agreeable

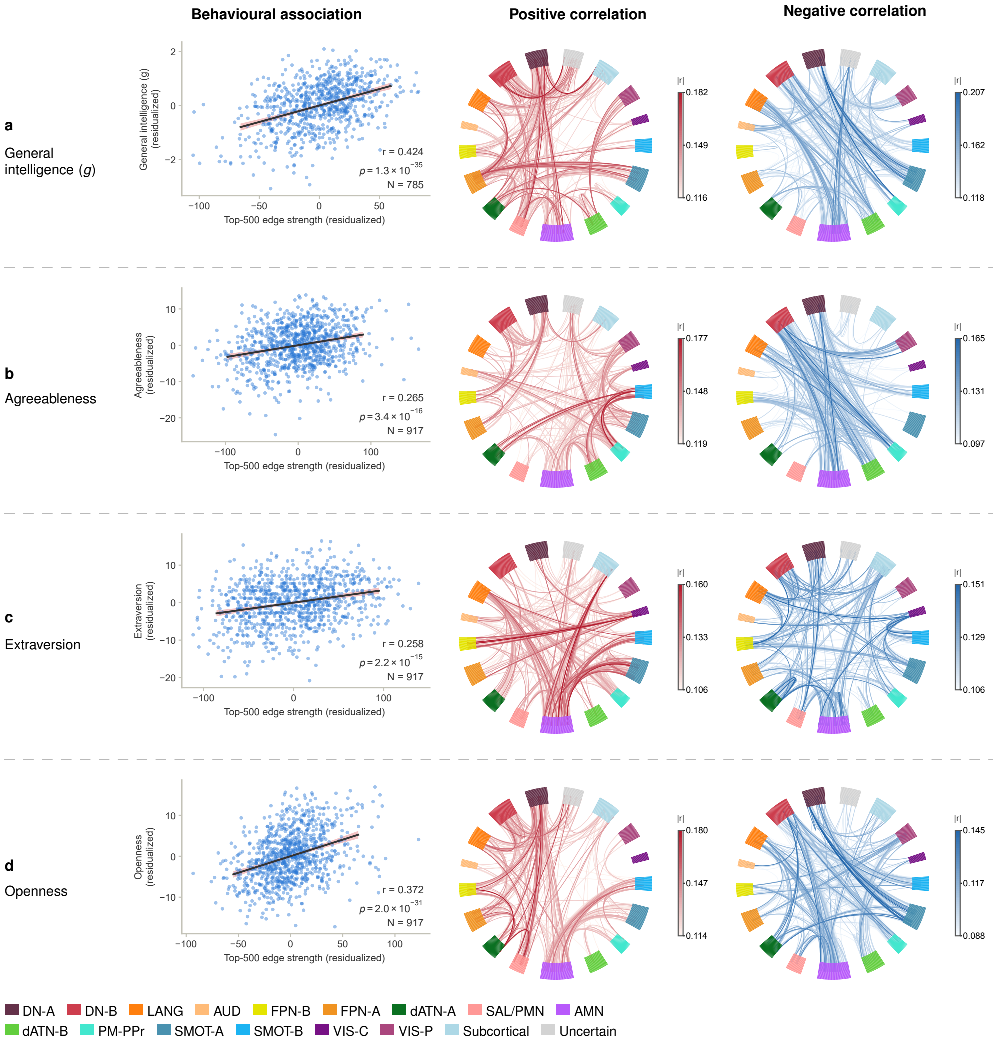


**Fig. B10 Whole-brain connectograms of the DMCF edges associated with intelligence and personality.** Each column displays one behavioral target: **a**, general intelligence $g$ ($N = 785$); **b**, agreeableness ($N = 917$); **c**, extraversion ($N = 917$); **d**, openness ($N = 917$). Left panels plot the residualized Top-500 edge strength (sum of the 500 positive minus the 500 negative edges) against the residualized behavioral score, with ordinary least-squares fits and 95% confidence bands. Middle and right panels show the whole-brain circular connectograms of the DMCF edges positively (red) and negatively (blue) correlated with each target across the 432 nodes (400 Schaefer cortical parcels ordered by the 15 Du-15 networks and 32 Tian subcortical nuclei; network colour legend at the bottom). Edge colour saturation scales with $|r|$ within each panel.

individuals combine cohesive internal self-integration with focused, goal-directed external attention, keeping the two domains functionally insulated. In contrast, extraversion was visually dominated by the integrative hub role of the AMN alongside notable connections with subcortical nuclei: positive edges radiated densely from the AMN to third-tier somatomotor cortices (SMOT-A/B), language regions, and subcortical structures while also bridging first-tier networks with second- and third-tier visual and sensorimotor systems, whereas negative edges fell primarily within the first and third tiers and between the AMN and default/subcortical regions. Such pronounced AMN-centred cross-tier and subcortical integration aligns with the heightened external action propensity and affective reactivity characteristic of extraverted individuals. Meanwhile, for openness, positive edges coupled first-tier (DN-A/B) with second-tier (SAL/PMN, AMN) networks and linked the AMN to third-tier somatomotor cortex, whereas negative edges segregated upper-tier default and action systems from second- and third-tier sensorimotor, language, and visual cortices. This configuration likely enables open individuals to integrate internal representations with imaginative processing while shielding internal states from external sensory disruption. Across both intelligence and personality, DMCF signatures thus capture a trait-specific balance between hierarchical integration and segregation across large-scale functional systems.

#### B.7 Subcortical degree maps and key-node connectograms

While the whole-brain connectograms of Section B.6 delineate which large-scale cortical systems are bridged by behaviorally relevant DMCF edges, they do not resolve the specific subcortical structures that anchor these associations. To localize the subcortical hubs driving each phenotype, we quantified nodal participation (degree) across the 32 Tian subcortical nuclei by counting the positive and negative behaviorally associated edges ($p < 0.05$) incident to each bilateral nucleus, and identified the nuclei with the highest positive and negative degrees as key seed nodes (Section A.22).

Figure B11 displays the resulting subcortical degree maps across four surface views alongside the subcortico-cortical connectograms of the positive key nodes, while the connectograms of the negative key nodes are presented in Fig. B12. Across all key nodes, the net nodal strength (sum of positive minus negative DMCF edges) correlated significantly with the corresponding behavioral score ($r = 0.125$–$0.181$, all $p \leq 1.6 \times 10^{-4}$; $N = 785$ for $g$ and 917 for the personality traits), confirming that the aggregate co-occurrence frequency of these subcortical seeds meaningfully tracks individual differences in cognition and personality.

General intelligence ($g$) and agreeableness converged on a common striato-thalamic axis: in both phenotypes, the posterior caudate (pCAU) emerged as the primary positive key node (Fig. B11**a**–**d**), whereas dorsal thalamic nuclei served as the negative key nodes—the dorsoposterior thalamus (THA-DP) for $g$ and the dorsal anterior thalamus (THA-DA) for agreeableness (Fig. B12**a**)[14]; their net nodal strength tracked the corresponding behavioural score ($r = 0.181$, $p = 3.3 \times 10^{-7}$ for pCAU–$g$; $r = 0.152$, $p = 4.0 \times 10^{-6}$ for pCAU–agreeableness; $r = 0.178$, $p = 5.4 \times 10^{-7}$ for THA-DP–$g$; $r = 0.146$, $p = 8.6 \times 10^{-6}$ for THA-DA–agreeableness). In $g$, the cortical projections of the pCAU were organized into two prominent bundles, operating as a functional switcher between the default-mode (DN) and action-mode (AMN) networks[16, 53, 54]: positive edges concentrated densely within the DN and language networks, whereas negative edges targeted the AMN and adjacent sensorimotor cortices (Fig. B11**a**). This dual motif indicates that higher intelligence is supported by frequent dynamic co-occurrence (information exchange) between the pCAU and default/language systems, paired with functional segregation between the pCAU and the AMN. Such pCAU–AMN decoupling may facilitate cognitive performance either by shielding action-oriented systems from striatal interference so they can remain task-focused, or by reflecting an alternative, anti-correlated mode of information exchange between default-related and action-mode circuitry, consistent with recent evidence of inverse

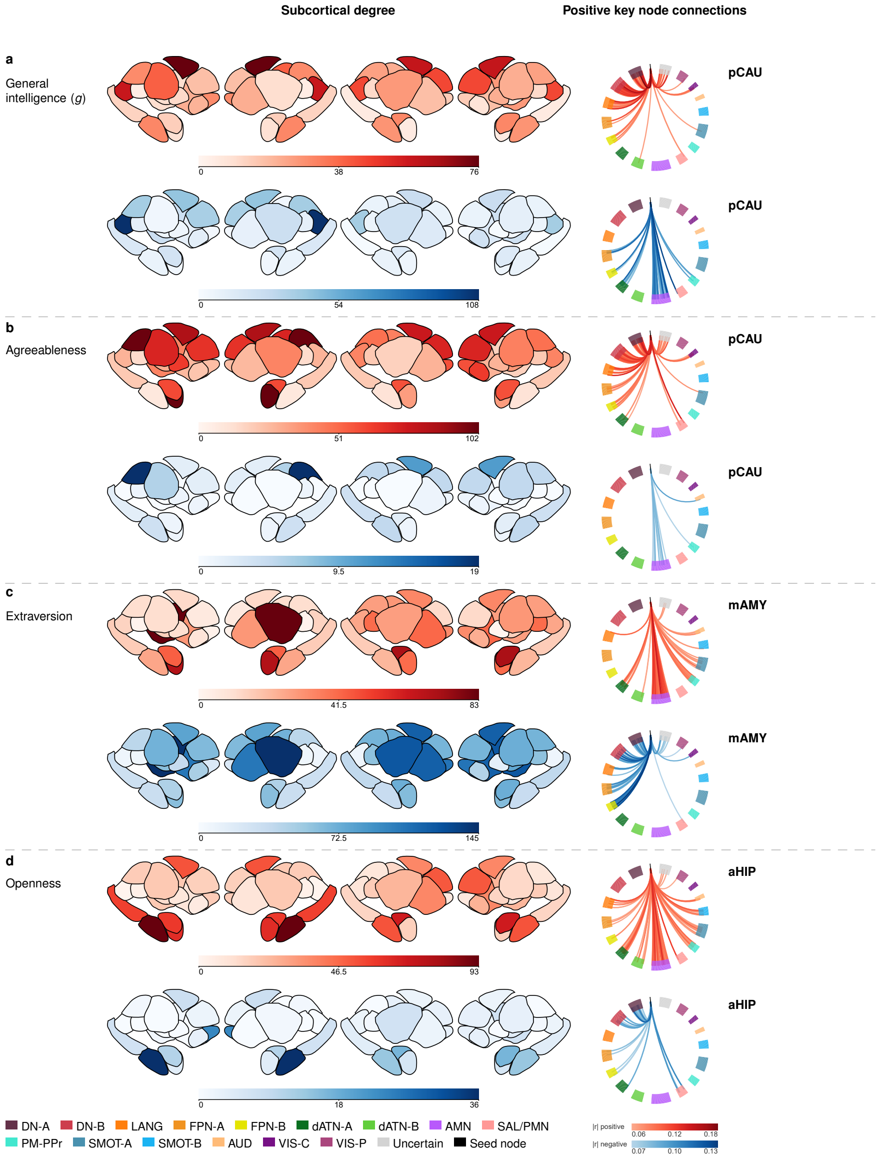


**Fig. B11 Subcortical degree maps and positive key-node connectograms.** Each target occupies two rows (upper, positive; lower, negative), with one letter per target, shown on the upper row: **a**, general intelligence $g$; **b**, agreeableness; **c**, extraversion; **d**, openness. Left, number of uncorrected $P < 0.05$ edges per nucleus of the Tian atlas (four views, both hemispheres; red, positive; blue, negative), each map on its own 0-to-maximum scale. Right, connections of the positive key node (highest positive count) in both rows—positive edges above, negative below—weighted by $|r|$, with one shared scale per sign; the two bilateral nuclei are black on top of the 400 Schaefer parcels (legend below). Node names are omitted. Negative key nodes: Fig. B12, except openness, whose positive and negative key node are the same nucleus.

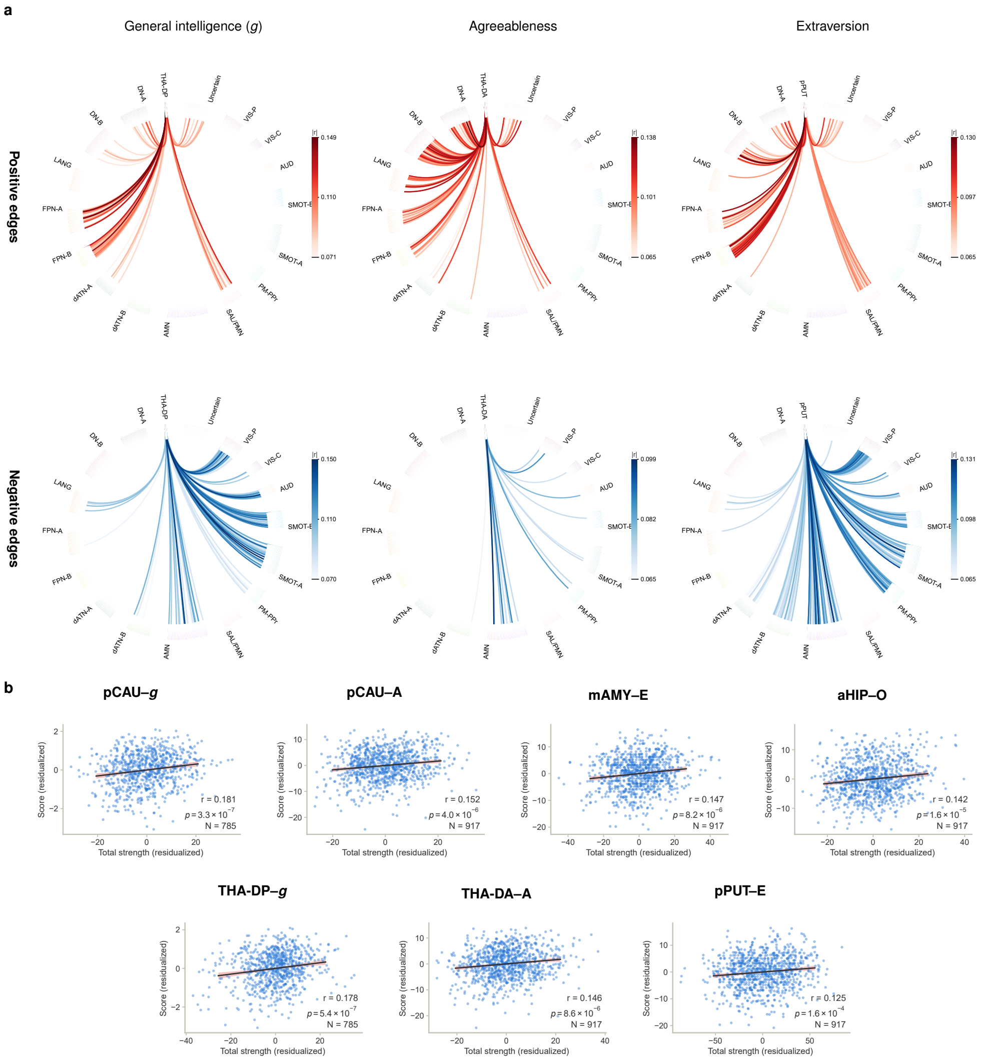


**Fig. B12 Connectograms of the negative key nodes and single-node associations. a**, Connectograms of the three negative key nodes, one per column: dorsoposterior thalamus (THA-DP) for general intelligence $g$; dorsal anterior thalamus (THA-DA) for agreeableness; posterior putamen (pPUT) for extraversion. Upper row, positive edges; lower row, negative edges; every plotted edge passes an uncorrected $P < 0.05$. Each panel shows the 402 nodes (two seeds, black; 400 Schaefer parcels) with the 16 network names round the ring and its own $|r|$ bar. **b**, Single-node associations: upper row, the positive key nodes pCAU–$g$, pCAU–agreeableness, mAMY–extraversion and aHIP–openness; lower row, the negative key nodes THA-DP–$g$, THA-DA–agreeableness and pPUT–extraversion. Points are participants; $x$, total node strength (summed positive minus summed negative DMCF weights); $y$, behavioural score; both residualized for age and sex. Lines, ordinary least-squares fits with 95% confidence bands. Openness has no ring because its positive and negative key node are the same nucleus (anterior hippocampus, aHIP; Fig. B11**d**); its association is shown in the scatter. All values are in-sample and descriptive.

BOLD signalling across these systems[126]. Strikingly, the pCAU exhibited a highly similar connectogram for agreeableness—dominated by positive coupling with the DN and negative coupling with the AMN (Fig. B11**b**). Because a participant's underlying DMCF matrix is invariant across behavioral tests, this convergence demonstrates that general intelligence (particularly its verbal component) and prosocial agreeableness share a common corticostriatal substrate centred on pCAU–DN integration. Quantitatively, 106 of the 203 pCAU edges associated with $g$ (52%) were also associated with agreeableness (60% of its 176 edges), and every shared edge carried the same sign (103 positive and 3 negative); moreover, 88 of these 106 shared edges (83%) were also among the 202 pCAU edges that tracked verbal intelligence (Factor 2), indicating that the shared pCAU substrate is anchored on the verbal component of $g$.

In contrast, extraversion and openness were anchored by socio-emotional, motor-gating, and mnemonic hubs that decoupled from first-tier regulatory networks while integrating with external action and perceptual systems. Extraversion inverted the striatal pattern observed in $g$ and agreeableness, recruiting the medial amygdala (mAMY) as its positive key node (Fig. B11**c**) and the posterior putamen (pPUT) as its negative key node (Fig. B12**a**), and standing out as the only target whose negative subcortical degree exceeded its positive degree. For the mAMY—a core hub for affective valuation and social feedback processing[58, 127]—positive edges coupled tightly with the AMN and sensorimotor networks, whereas negative edges segregated it from the first-tier DN (Fig. B11**c**)[59]; its net nodal strength was $r = 0.147$ ($p = 8.2 \times 10^{-6}$) for extraversion. This profile suggests that extraverted individuals anchor their affective responses in direct external action and sensory engagement rather than internal self-referential processing. Complementing the mAMY, the pPUT functioned as an executive motor gate for social action[55, 56]: its tighter integration with first-tier regulatory networks and stronger segregation from second- and third-tier attentional and sensorimotor networks (Fig. B12**a**; net nodal strength $r = 0.125$, $p = 1.6 \times 10^{-4}$) indicate that extraverts translate internal intentions into overt social behaviors with reduced motor inhibition[57]. Finally, openness was the unique phenotype in which a single structure—the anterior hippocampus (aHIP)—served as both the positive and negative key node (Fig. B11**d**). Consistent with the established role of the hippocampus in episodic memory and future scenario construction[60, 128], positive edges of the aHIP projected extensively to second- and third-tier attentional and sensorimotor networks, whereas negative edges concentrated within first-tier transmodal networks[61] (net nodal strength $r = 0.142$, $p = 1.6 \times 10^{-5}$). Greater co-occurrence with lower- and middle-tier networks likely enables open individuals to incorporate rich external sensory material to fuel constructive imagination, while reduced co-occurrence with first-tier networks reflects relaxed top-down executive control and disinhibition over imaginative states.

#### B.8 Factor-level decomposition of the key-node associations

To resolve the composite trait signatures of Section B.7 into more unitary behavioral dimensions, we projected the subcortico-cortical associations of the pCAU, mAMY, and aHIP onto their related latent factors (Section A.23; Fig. B13). At the pCAU, the composite $g$ connectogram cleanly dissociated into two complementary cognitive axes (Fig. B13**a**): verbal intelligence (Factor 2) selectively captured the positive bundle—frequent co-occurrence with first-tier default and language networks (DN-A/B and LANG) supporting semantic and verbal reasoning—whereas visual intelligence (Factor 8) captured the negative bundle, where decoupling from action and second-/third-tier sensorimotor networks (AMN, dATN-A/B, PM-PPr, SMOT-A/B, and VIS-C/P) likely shields lower-tier perceptual systems from striatal interference to facilitate visual-spatial processing. Notably, comparing this decomposition with the agreeableness profile in Fig. B11**b**—for which the pCAU also served as the primary positive key node—revealed a striking resemblance between the connectograms of agreeableness and verbal intelligence, indicating that prosocial agreeableness shares a common corticostriatal substrate with verbal cognitive ability.

At the mAMY, all three socio-affective factors shared a consistent limbic–action motif anchored by tighter coupling with the AMN and SMOT-A/B and decoupling from DN-A/B (Fig. B13**b**). While social distress (Factor 1) formed a direct mirror image of extraversion, well-being (Factor 4) and especially extraversion and friendship (Factor 5) closely recapitulated the whole-trait ring, confirming that mAMY–AMN co-occurrence directly supports external social engagement and friendship formation[58, 127]. Finally, at the aHIP, processing speed (Factor 3) and visual intelligence (Factor 8) engaged the same anatomical connections across first-tier transmodal (DN-A/B and FPN-A/B) and second-/third-tier perceptual-motor networks (dATN-A/B, AMN, SMOT-A/B, and VIS-C/P), but with opposite polarity (Fig. B13**c**): stronger aHIP coupling with second-/third-tier networks and decoupling from first-tier networks—the hallmark pattern of trait openness—predicted higher visual intelligence but slower processing speed. Thus, highly open individuals exhibit a neural component characterized by strong visual-spatial construction alongside slower processing speed, matching their behavioral tendency toward broad sensory intake and unhurried imaginative exploration[60, 61, 128]. Taken together, these factor-level decompositions demonstrate that broad personality traits are not purely affective constructs, but encompass both socio-emotional dimensions and distinct cognitive-intellectual components in their underlying brain dynamics.

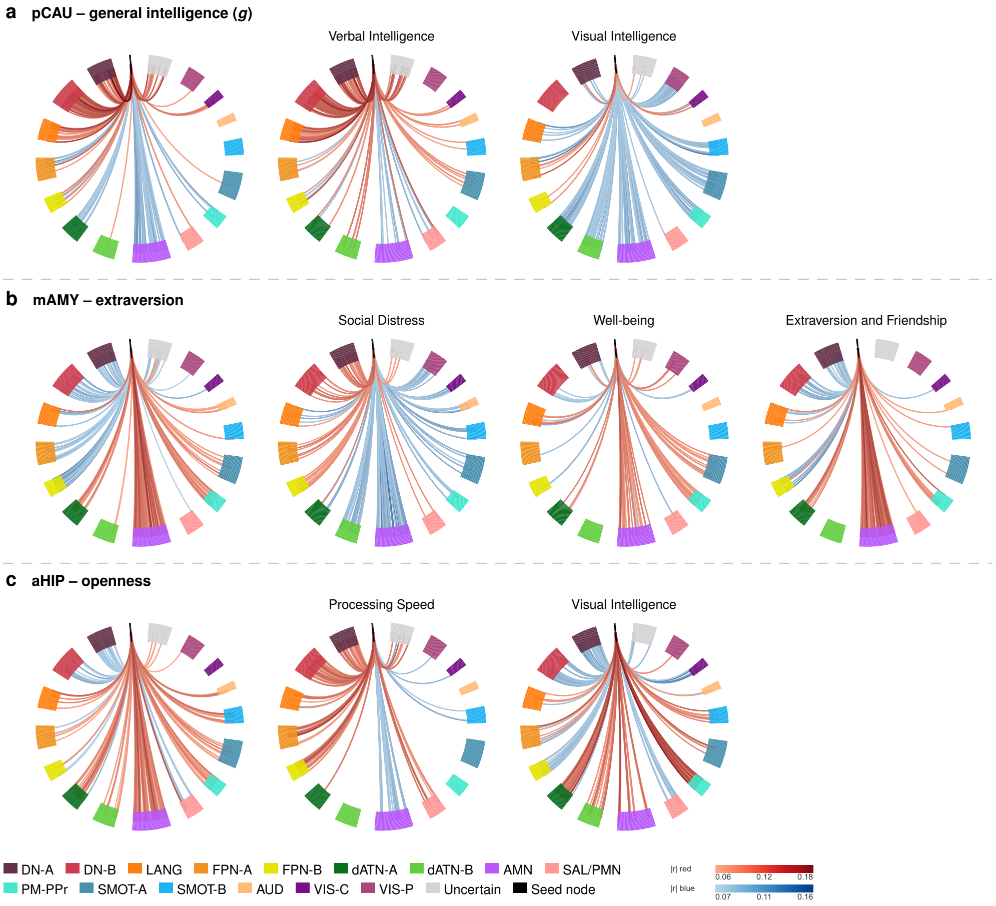


**Fig. B13 Factor-level decomposition of the key-node associations. a–c**, Subcortico-cortical connectograms ($p < 0.05$, ordered by the Du-15 networks) for each seed node with its broad trait (Column 1) and related latent factors ($N = 779$; subsequent columns): **a**, pCAU with $g$ ($N = 785$) and Factors 2 and 8; **b**, mAMY with extraversion ($N = 917$) and Factors 1, 4, and 5; **c**, aHIP with openness ($N = 917$) and Factors 3 and 8. Red denotes connections associated with higher trait scores, higher cognition (Factors 2, 3, and 8), or higher extraversion/friendship (Factors 4 and 5), whereas Factor 1 is plotted in its native social-distress direction. Colour saturation encodes $|r|$ on a shared scale per sign.

#### B.9 Cross-cultural validation and functional dissociation in the ISYB cohort

To test whether the subcortico-cortical dynamic architecture identified in HCP generalizes across populations, we applied the identical analytical framework to the East Asian ISYB cohort (Section A.24). Lacking the extensive cognitive battery used for HCP, we extracted a composite cognitive dimension via maximum-likelihood factor analysis of six age- and sex-residualized behavioral indices ($N = 165$; Fig. B14). This yielded a target cognitive factor (Factor 2, capturing visuospatial processing speed and conscientiousness) which we evaluated alongside NEO agreeableness, extraversion, and openness. At the whole-brain level, the top 500 positive and negative DMCF edges remained strongly correlated with all four behavioral targets in ISYB ($r = 0.615$–$0.804$; Fig. B15), though specific connectogram topologies shifted, reflecting both cross-cultural divergence and higher sampling variability in the smaller cohort.

Focusing on the 32 subcortical nuclei, we observed shifts in exact nodal degree rankings ($p < 0.05$ edges; Fig. B16). Most notably, the dominant hub for openness shifted along the hippocampal longitudinal axis[118, 129] from the anterior hippocampus (aHIP) in HCP to the posterior hippocampus (pHIP) in ISYB, potentially reflecting a cultural-to-physiological adaptation from Western gist-based divergent imagination to East Asian context-rich memory elaboration[130, 131]. Despite these shifts, all seven HCP-defined primary key nodes (pCAU, THA-DP, THA-DA, mAMY, pPUT, and aHIP) retained highly significant within-sample associations with their corresponding phenotypes in the Chinese cohort ($r = 0.322$–$0.463$, all $p < 10^{-4}$). This suggests that while overarching subcortical systems show broad cross-population relevance, their localized subnuclear recruitment is sensitive to cultural context.

Examining the subcortico-cortical connectograms of these replicated key nodes in ISYB (Fig. B17) revealed a clear cross-cultural principle: the routing topology of cognitive computation was conserved, whereas the representations of socio-emotional personality diverged. In the cognitive and experiential domains, key nodes maintained consistent cross-hierarchy routing motifs across cohorts. Specifically, pCAU and THA-DP supported Factor 2 cognitive performance through conserved interactions with the action-mode (AMN) and dorsal attention (dATN) systems (Fig. B17**a**), paralleling the role of pCAU in HCP as a functional switcher between default-mode and action-mode networks. Similarly, for openness, the aHIP replicated the HCP pattern of reduced co-occurrence with first-tier regulatory networks alongside greater integration with second- and third-tier communities (Fig. B17**d**), reflecting how open individuals incorporate broader external sensory material to fuel constructive imagination under reduced top-down inhibition[132, 133]. By contrast, socio-emotional routing showed strong cultural specificity (Fig. B17**b,c**), most notably in the mAMY network for extraversion, where most shared significant edges exhibited opposite signs between cohorts (Fig. B17**c**). In the Western HCP cohort, high extraversion was driven by tighter integration of the mAMY with the AMN and sensorimotor networks alongside decoupling from first-tier networks, reflecting active external seeking and direct action engagement[134]. In the East Asian ISYB cohort, however, high extraversion relied on mAMY segregation from the dorsal attention networks (dATN-A and dATN-B)—a release from internal cognitive and attentional monitoring that aligns with collectivist cultural norms of behavioral restraint[64, 135].

To directly test the hypothesis that core cognitive routing forms a cross-culturally conserved computational backbone whereas socio-emotional circuits tailor their routing strategies to cultural ecologies, we conducted a three-part cross-dataset validation (Fig. B18). First, we transferred the intelligence-related edges of pCAU and THA-DP identified in ISYB (Factor 2) directly to the Western HCP cohort without re-estimation. Both ISYB-defined subcortical networks significantly tracked HCP general intelligence $g$ out-of-sample (pCAU: $r = 0.080$, $p = 0.025$; THA-DP: $r = 0.097$,

$p = 0.006$; $N = 785$; Fig. B18**a**), confirming that the dynamic subcortical routing underlying core cognitive computation generalizes across populations. Second, to test whether the cross-culturally conserved aspects of personality traits are specifically their embedded cognitive components, we extracted the cross-dataset consistent (same-sign intersection) edges between ISYB and HCP for the pCAU agreeableness network and the aHIP openness network. In line with our hypothesis, the cross-culturally conserved pCAU agreeableness network significantly predicted general intelligence $g$ in HCP ($r = 0.104$, $p = 0.0035$, $N = 785$; Fig. B18**b**), reflecting a shared substrate for language-based cognition and pro-social conformity, whereas the conserved aHIP openness network specifically tracked HCP visual intelligence (Factor 8; $r = -0.102$, $p = 0.0046$, $N = 779$; Fig. B18**c**).

Third, we examined how the culturally divergent mAMY pathways support social affiliation within each cultural ecology. Despite the striking cross-cultural divergence in specific cortical pathways between East Asian and Western extraversion, extraversion and agreeableness converged within each cohort onto a shared mAMY routing substrate (the within-cohort intersection of mAMY edges across the two traits). Crucially, this shared mAMY substrate significantly predicted perceived social support in ISYB (MSPSS; $r = 0.229$, $p = 0.0034$, $N = 162$; Fig. B18**d**) and the extraverted friendship factor in HCP ($r = 0.094$, $p = 0.0045$, $N = 916$; Fig. B18**e**), while relying on distinct cortical connections in the two populations (compare connectograms in Fig. B18**d** and **e**). Together, these cross-dataset analyses demonstrate that whereas cognitive routing through the caudate, thalamus, and hippocampus provides a universal, cross-culturally invariant computational backbone for both intelligence and the cognitive dimensions of personality, subcortical socio-emotional circuits through the medial amygdala dynamically adapt their cortical routing strategies to local cultural norms while converging functionally to support human social affiliation[136, 137].

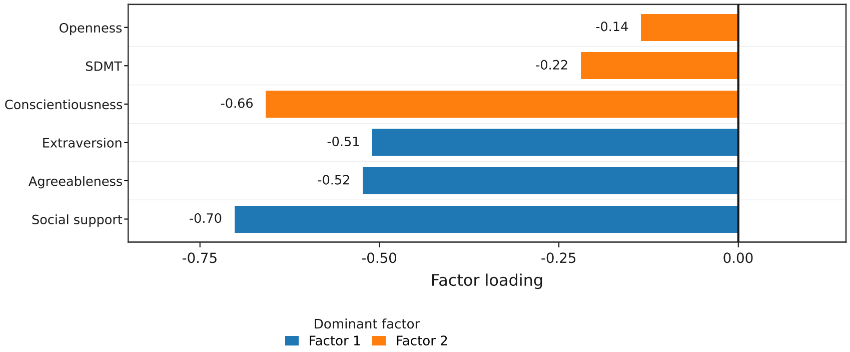


**Fig. B14 Factor loadings of the ISYB behavioral structure.** Maximum-likelihood factor analysis with varimax rotation across six age- and sex-residualized behavioral indices ($N = 165$). Each index is plotted on its dominant factor: Factor 1 (blue; social support, agreeableness, and extraversion) and Factor 2 (orange; SDMT cognitive performance and conscientiousness). Loadings are uninverted, such that lower factor scores indicate higher performance or trait levels.

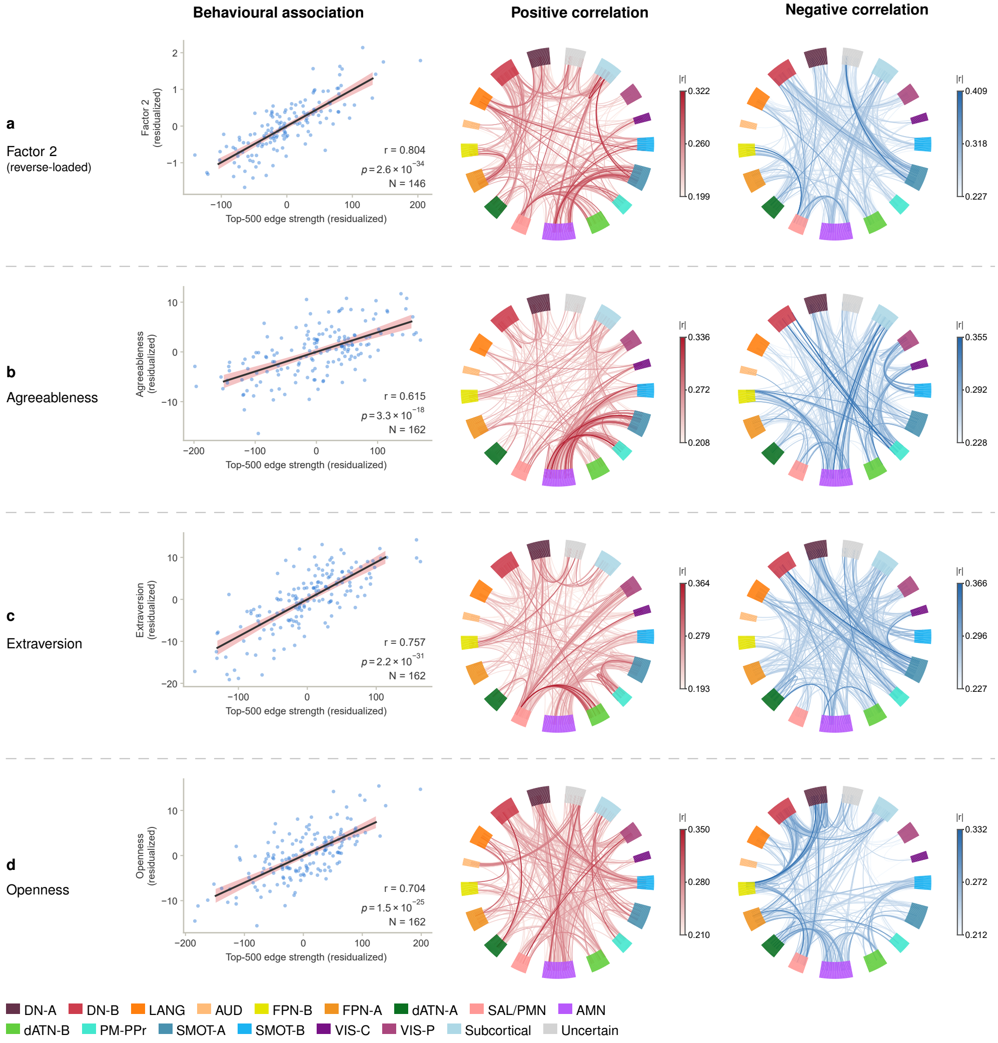


**Fig. B15 Whole-brain DMCF connectograms in ISYB. a**, Factor 2 ($N = 146$; signs inverted so red denotes higher cognitive performance). **b**, Agreeableness ($N = 162$). **c**, Extraversion ($N = 162$). **d**, Openness ($N = 162$). Left, residualized Top-500 net co-occurrence frequency versus behavioral scores (linear fits with 95% confidence bands). Middle and right, 432-node connectograms (400 Schaefer parcels and 32 Tian nuclei) of positively (red) and negatively (blue) correlated edges, with saturation scaled by $|r|$.

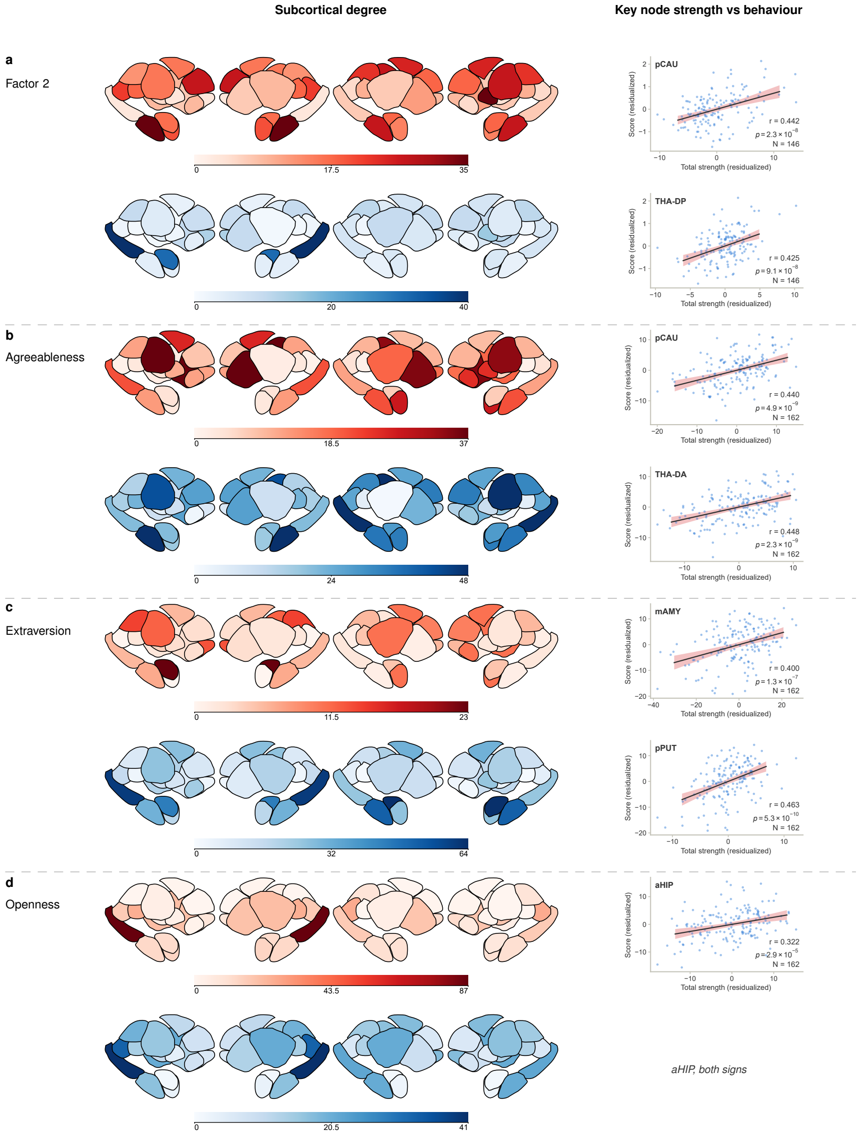


ISYB, one resting-state run. Left: number of incident edges with uncorrected $p < 0.05$, each map on its own 0-to-maximum scale.
Right: total node strength (sum of positive minus negative edge strengths) against the behavioural score,
both residualised for age and sex; coefficients are in-sample.

**Fig. B16 Subcortical degree maps and key-node replication in ISYB.** **a**, Factor 2. **b**, Agreeableness. **c**, Extraversion. **d**, Openness. Left, nodal degree of $p < 0.05$ positive (red, upper row) and negative (blue, lower row) edges across the Tian subcortical atlas. Right, associations between residualized net nodal frequency and behavioral scores for the seven HCP-defined key nodes (Section B.7; aHIP serves as both positive and negative hub for openness).

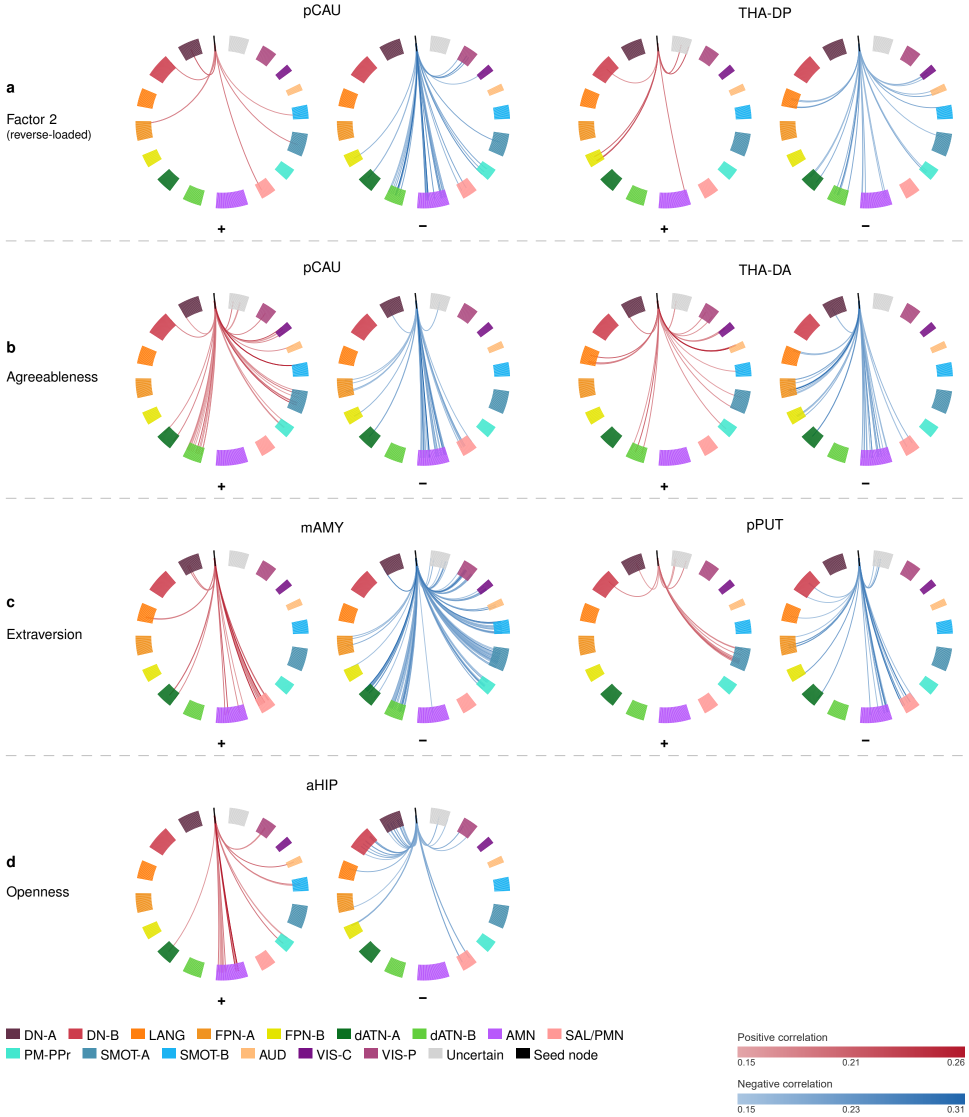


**Fig. B17 Subcortico-cortical connectograms of the key nodes in ISYB.** Rows **a**–**d** display the positively (left rings, red) and negatively (right rings, blue) correlated subcortico-cortical DMCF edges ($p < 0.05$) for each prior key node across the four behavioral targets: **a**, pCAU and THA-DP for Factor 2; **b**, pCAU and THA-DA for agreeableness; **c**, mAMY and pPUT for extraversion; **d**, aHIP for openness (serving as both positive and negative hub). Each ring comprises 402 nodes (two bilateral subcortical seeds in black and 400 Schaefer cortical parcels ordered by the Du-15 networks), with edge saturation scaled by $|r|$. In **a**, ring polarity is inverted so that red consistently indicates higher cognitive performance.

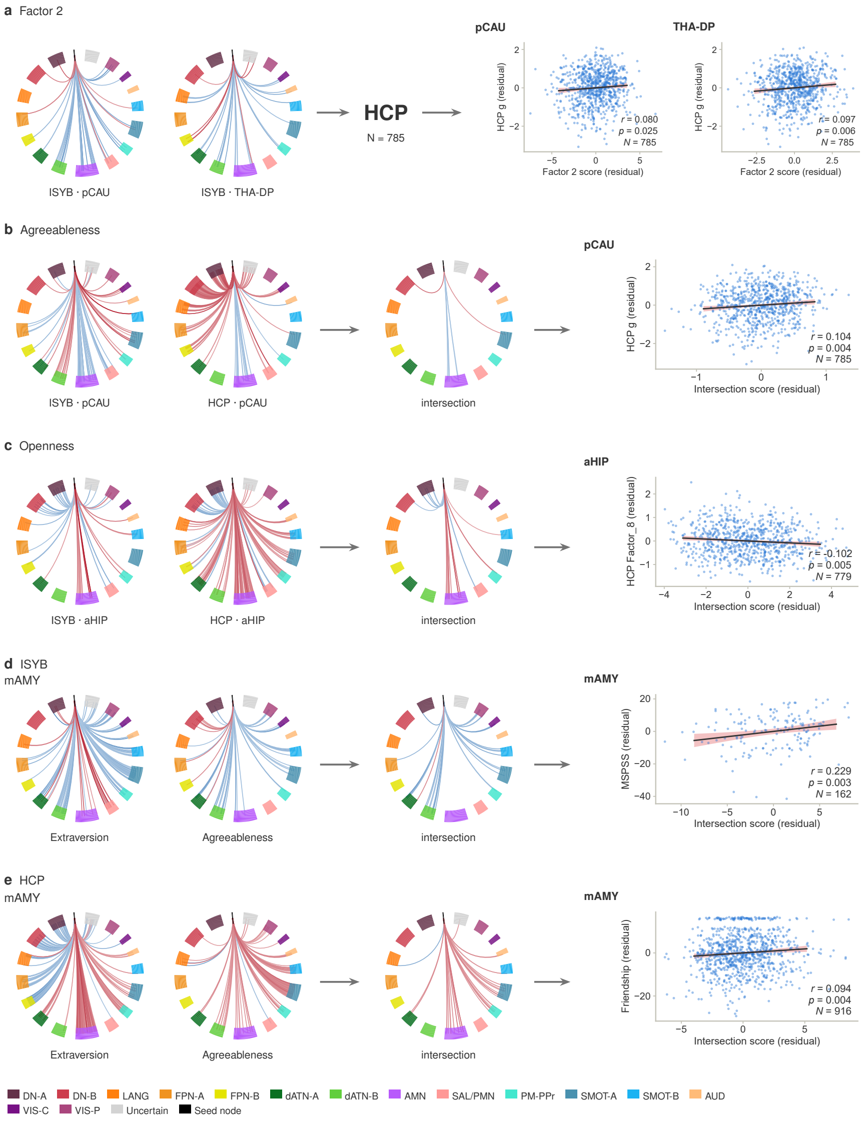


**Fig. B18 Cross-dataset validation of the conserved cognitive backbone and culturally adapted socio-emotional routing.** **a**, Out-of-sample transfer of ISYB Factor 2 edges (pCAU and THA-DP; signs inverted) predicting HCP $g$ ($N = 785$). **b**,**c**, Cross-dataset consistent edges (ISYB–HCP same-sign intersection) of the pCAU agreeableness network predicting HCP $g$ ($N = 785$; **b**) and of the aHIP openness network tracking HCP visual intelligence (Factor 8, $N = 779$; **c**). **d**,**e**, Within-cohort mAMY convergence (agreeableness–extraversion intersection) predicting perceived social support in ISYB ($N = 162$; **d**) and friendship scores in HCP ($N = 916$; **e**).
